# Profibrotic monocyte-derived alveolar macrophages as a biomarker and therapeutic target in systemic sclerosis-associated interstitial lung disease

**DOI:** 10.1101/2025.08.07.669006

**Authors:** Nikolay S. Markov, Anthony J. Esposito, Karolina J. Senkow, Maxwell Schleck, Luisa Cusick, Zhan Yu, Yuliana V. Sokolenko, Estefani Diaz, Emmy Jonasson, Suchitra Swaminathan, Ziyan Lu, Radmila Nafikova, Samuel Fenske, Elsie G. Bunyan, Xóchitl G. Pérez-Leonor, Hiam Abdala-Valencia, Annette S. Flozak, Nikita Joshi, A. Christine Argento, Elizabeth S. Malsin, Paul A. Reyfman, Jonathan Puchalski, Mridu Gulati, Mary Carns, Kathleen Aren, Phillip Cooper, Natania S. Field, Suror Mohsin, Malek Shawabkeh, Alexandra Soriano, Aaron N. Gundersheimer, Isaac A. Goldberg, Bailey Damore, Alec Peltekian, Ankit Agrawal, Crystal Cheung, Stephanie Perez, Shannon Teaw, Alyssa Williams, Nicolas Page, Sophia E. Kujawski, William Odell, Baran Ilayda Gunes, Michelle Cheng, Morgan Emokpae, R. Ian Cumming, Robert M. Tighe, Kevin Grudzinski, Hatice Savas, Ami N. Rubinowitz, Bashar A. Kadhim, Chitaru Kurihara, Ankit Bharat, Vikas Mehta, Jane E. Dematte, Bradford C. Bemiss, Hadijat M. Makinde, Carla M. Cuda, Matthew Dapas, Carrie Richardson, Harris Perlman, Anna P. Lam, Cara J. Gottardi, G.R. Scott Budinger, Alexander V. Misharin, Monique E. Hinchcliff

## Abstract

Interstitial lung disease (ILD) is present in over 60% of patients with systemic sclerosis (SSc) and is the leading cause of SSc-related deaths. Profibrotic monocyte-derived alveolar macrophages (MoAM) play a causal role in the pathogenesis of pulmonary fibrosis in animal models where their persistence in the niche requires signaling through Colony Stimulating Factor 1 Receptor (CSF1R). We hypothesized that the presence and proportion of MoAM in bronchoalveolar lavage (BAL) fluid from patients with SSc-ILD may be a biomarker of ILD severity. To test this hypothesis, we analyzed BAL fluid from 9 prospectively enrolled patients with SSc-ILD and 13 healthy controls using flow cytometry and single-cell RNA sequencing. Patients with SSc-ILD had more MoAM and interstitial macrophages in BAL fluid than healthy controls, and their abundance was associated with lung fibrosis severity. We identified changes in the MoAM transcriptome as a function of treatment with mycophenolate, an effective therapy for SSc-ILD. In SSc-ILD lung explants, spatial transcriptomics identified an expanded population of interstitial macrophages spilling into the alveolar space. Our findings suggest that the proportion of profibrotic MoAM and interstitial macrophages in BAL fluid may serve as a biomarker of SSc-ILD and credential them as possible targets for therapy.

## Introduction

Systemic sclerosis (SSc) is an autoimmune rheumatic disease characterized by tissue inflammation followed by progressive fibrosis in multiple organs. Interstitial lung disease (ILD) affects over 60% of patients with SSc and is the leading cause of SSc-associated death or lung transplantation^1,2^. In most patients, SSc-ILD progresses slowly; however, up to a quarter will rapidly progress to respiratory failure^3^. Therapies available to patients with SSc-ILD slow disease progression but do not reverse established fibrosis. Many of these agents, including mycophenolate mofetil, cyclophosphamide, tocilizumab, and rituximab target inflammatory pathways^4^.

We and others have shown that a population of profibrotic monocyte-derived alveolar macrophages (MoAM) is necessary for lung fibrosis induced by bleomycin, asbestos, or radiation in mice^5–10^ where their maintenance in the lung requires signaling through colony stimulating factor-1/colony stimulating factor-1 receptor (CSF1/CSF1R). These findings were recently credentialed in patients with refractory multiple organ fibrosis resulting from chronic graft-versus-host disease after allogeneic stem cell transplantation, where an inhibitory antibody targeting the CSF1R, axatilimab, reversed established fibrosis across multiple organs, including the lung and skin, leading to its FDA approval^11^. Importantly, we and others have identified a population of macrophages with a transcriptional signature resembling murine MoAM in lung explants from patients with pulmonary fibrosis including SSc-ILD^12–17^. Moreover, transcriptionally similar populations of monocyte-derived macrophages have been implicated in fibrosis in other organs^18,19^.

Alveolar macrophages can be safely and serially obtained from bronchoalveolar lavage (BAL) fluid and analyzed using flow cytometry or genomic assays^20,21^. In this study, we show the abundance of a specific alveolar macrophage subset—profibrotic MoAM—is increased in BAL fluid from patients with SSc-ILD compared to healthy controls and is associated with SSc-ILD severity measured by pulmonary function testing and fibrosis severity on chest high-resolution computed tomography. Single-cell RNA-sequencing analysis identified both shared and cell type-specific changes in gene expression associated with SSc-ILD and treatment with mycophenolate. We found that expression of CSF1R was increased in precursors of profibrotic MoAM. We used single-cell spatial transcriptomics to localize MoAM to the alveolar space in lung explants from patients with SSc-ILD. We also identified an expansion of interstitial macrophages in patients with SSc-ILD. These interstitial macrophages were associated with a multicellular niche enriched in T cells and were spilling into the alveolar space. Together, these findings support the use of BAL sampling to measure the abundance and transcriptional profile of MoAM and interstitial macrophages as a biomarker and possible therapeutic target for patients with SSc-ILD.

## Results

### Study cohort

Nine patients with SSc-ILD (patients) and 13 healthy volunteers (controls) were recruited and underwent bronchoscopy for research purposes (**Table 1**). The majority of both groups were white and biologically female and differed only by age (mean age 54 vs. 29 years, respectively). Four and five patients were classified as having limited cutaneous and diffuse cutaneous SSc, respectively. At the time of bronchoscopy the median time since SSc diagnosis was 16.9 years (IQR 8.7–19.4), and median time since radiographically confirmed SSc-ILD diagnosis was 8.1 years (IQR 4.3–11.7). The median modified Rodnan skin score (mRSS) was 6 (IQR 4–11). Five patients had positive Scl-70 serum autoantibodies. Pulmonary symptoms (cough and/or dyspnea) were reported by seven patients. Pulmonary function test data revealed that all nine patients had restrictive lung physiology with median (IQR) % predicted forced vital capacity (FVC) 70 (65.9–74.4) and % predicted diffusion capacity for carbon monoxide (DLCO) 55.7 (48.7–62.8). Average Kazerooni scores in all five lung lobes were 1.6 (1.2–2.4) for ground glass opacity (inflammation) and 2.0 (1.2–2.4) for fibrosis, both higher in the lower lobes compared to upper lobes (**Table 2**).

**Table 1:** Patient and control characteristics at the time of bronchoscopy. SSc-specific antibodies test, C-reactive protein, blood monocytes, modified Rodnan skin score, pulmonary function tests, 6-minute walking tests, right heart catheterization, and HRCT were used from dates closest to the bronchoscopy. For timing of PFT, HRCT and therapy, see **Figure S1a**. Within-group values are reported as medians and 1st and 3rd quartiles. When data is missing, the effective n is indicated for the measurement within the group. Abbreviations: dcSSc: diffuse cutaneous SSc; lcSSc: limited cutaneous SSc; FVC: forced vital capacity; FEV1: forced expiratory volume in 1 second; TLC: total lung capacity; DLCO: diffusing capacity for carbon monoxide; RVSP: right ventricular systolic pressure; 6MWD: 6-minute walking distance; HRCT: high-resolution computed tomography.

|  |  | SSc | control |
| --- | --- | --- | --- |
| n |  | 9 | 13 |
| Age, years, median [Q1,Q3] |  | 53.8 [37.9,60.4] | 29.0 [24.0,34.0] |
| Sex, n | Female | 6 (67%) | 9 (69%) |
|  | Male | 3 (33%) | 4 (31%) |
| Race, n | African American | 1 (11%) | 2 (15%) |
|  | Caucasian | 5 (56%) | 8 (62%) |
|  | Hispanic or Latino | 3 (33%) | 2 (15%) |
|  | Asian | — | 1 (8%) |
| Smoker, n | current or former | 1 (11%) | — |
|  | never | 8 (89%) | 13 (100%) |
| SSc subtype, n | dcSSc | 5 (56%) | — |
|  | lcSSc | 4 (44%) | — |
| Time since diagnosis, years, median [Q1,Q3] | SSc | 16.9 [8.7,19.4] | — |
|  | SSc-ILD | 8.1 [4.3,11.7] | — |
| Symptoms, n | Cough | 6 (67%) | — |
|  | Dyspnea | 7 (78%) | — |
| SSc-specific autoantibodies, n | Anti-nuclear | 9 (100%) | — |
|  | Anti-topoisomerase I (Scl-70) | 5 (56%) | — |
|  | Anti-centromere | 0 (0%) | — |
|  | Anti-RNA polymerase III | 2 (25%)<br>(n = 8) | — |
| C-reactive protein, mg/l, median [Q1,Q3] |  | 6.8 [2.1,12.8]<br>(n = 8) | — |
| Blood monocyte %, median [Q1,Q3] |  | 8.0 [6.0,10.0] | — |
| Modified Rodnan skin score, median [Q1,Q3] | total | 6 [4,11] | — |
|  | lcSSc | 4 [3,6] | — |
|  | dcSSc | 8 [6,13] | — |
| Digital ulcer or pitting scars, n | digital tip ulcers | 3 (33%) | — |
|  | fingertip pitting scars | 3 (33%) | — |
| Medications, n |  | 4 (44%) | — |
|  | Mycophenolate | 4 (44%) | — |
|  | Rituximab | 1 (11%) | — |
| FVC % predicted, median [Q1,Q3] |  | 70.0 [65.9,74.4] | 108.0 [100.2,110.2]<br>(n = 8) |
| FEV1 % predicted, median [Q1,Q3] |  | 72.1 [64.4,76.1] | 95.5 [90.5,102.8]<br>(n = 8) |
| FEV1/FVC ratio, %, median [Q1,Q3] |  | 80.4 [78.3,84.7] | — |
| TLC % predicted, median [Q1,Q3] |  | 59.1 [53.7,62.6]<br>(n = 8) | — |
| DLCO % predicted, median [Q1,Q3] |  | 55.7 [48.7,62.8]<br>(n = 7) | — |
| Estimated RVSP, mmHg, median [Q1,Q3] |  | 27.0 [18.0,37.0]<br>(n = 8) | — |
| 6MWD distance, m, median [Q1,Q3] |  | 450.0 [432.0,456.0]<br>(n = 7) | — |
| Average HRCT score per lobe, median [Q1,Q3] | Ground glass opacity | 1.6 [1.2,2.4] | — |
|  | Fibrosis | 2.0 [1.2,2.4] | — |
| HRCT score in lavaged lobe, median [Q1,Q3] | Ground glass opacity | 2.0 [1.0,3.0] | — |
|  | Fibrosis | 2.0 [1.0,3.0] | — |

**Table 2:**
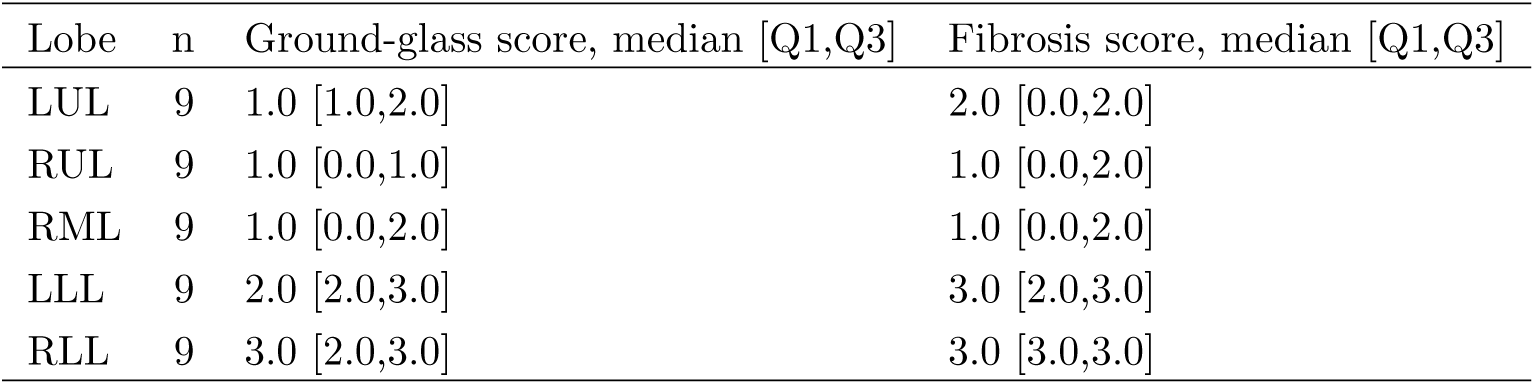
Summary of Kazerooni scores of patient HRCT scans from Table 1.

Four patients were prescribed immunosuppressive therapy at the time of bronchoscopy, three mycophenolate mofetil and one mycophenolic acid (**Figure S1a**). These four subjects were assessed by the treating clinician to be (1) stable on mycophenolate, (2) worsening on mycophenolate with dose escalation immediately prior to bronchoscopy, (3) stable on mycophenolate combination therapy with rituximab, and (4) stable on mycophenolate therapy following autologous stem cell transplant for SSc-ILD. No subjects had exposure to antifibrotic or supplemental oxygen therapy. Of the five subjects not taking immunosuppression, two were never treated due to assessment of stable ILD. The other three were previously treated, two with mycophenolate and one with methotrexate but had discontinued treatment due to stable, long-standing SSc-ILD (**Figure S1b**).

### Profibrotic MoAM are expanded in patients with SSc-ILD

We performed single-cell RNA-sequencing (scRNA-seq) on cells isolated from the BAL fluid from patients and controls. We resolved DCs, T cells, B cells, plasma cells, mast cells, and oral mucosa and airway epithelial cells, and multiple macrophage subsets (**Figure 1a**, **Figure S1c**, **Table S1**). We did not observe batch effects related to scRNA-seq chemistry, sample processing protocol, or study recruitment site (**Figure S1d**). Macrophage clusters (expressing *C1QA*, *MRC1*, *MSR1*) included mature tissue-resident alveolar macrophages (TRAM), characterized by expression of *FABP4*, *NUPR1*, and *INHBA*, and MoAM, characterized by the lack of FABP4 expression and expression of *VCAN* and *CCL2* (**Figure 1b**). Similar to previous reports^22–24^, we detected a cluster of macrophages expressing *SEPP1* (*SELENOP*), *CCL13*, and *FOLR2* that matched the expression profile of interstitial macrophages (**Figure 1b**).

**Figure 1:**
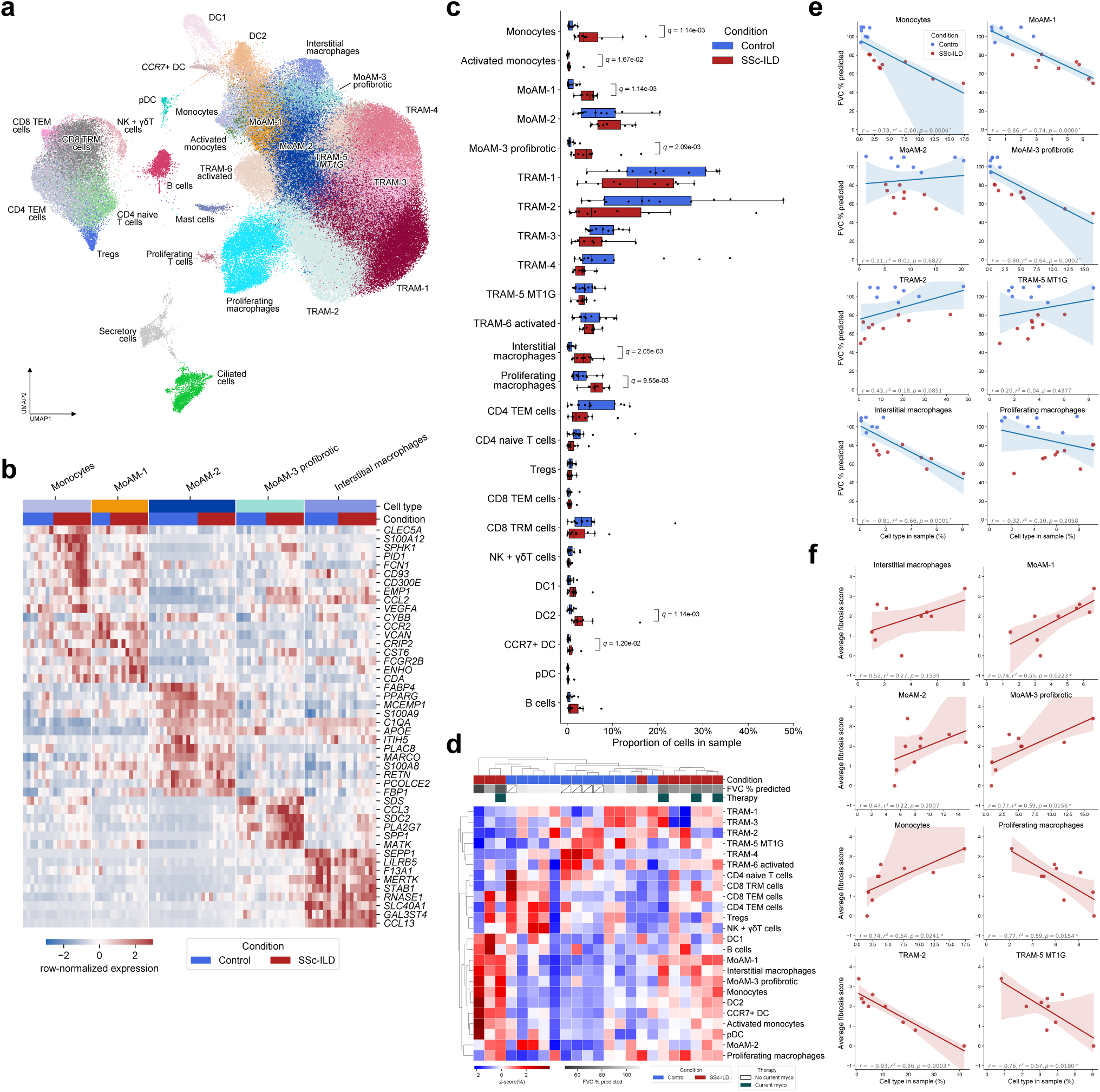
The abundance of profibrotic monocyte-derived alveolar macrophages in BALF is associated with worse lung function in patients with SSc-ILD. **a**. Uniform manifold approximation and projection (UMAP) plot showing integrated analysis of BAL immune cells from patients with SSc-ILD (*n* = 9) and healthy volunteers (*n* = 13). Tissue-resident alveolar macrophages (TRAM), monocyte-derived alveolar macrophages (MoAM), type I conventional DCs (DC1), type II conventional DCs (DC2), migratory dendritic cells (CCR7+ DC) and plasmacytoid dendritic cells (pDC). **b**. Heatmap of expression of macrophage subset-specific genes. **c**. Proportions of cell clusters represented in the UMAP in a, Mann-Whitney U tests with FDR correction. *q*-values < 0.05 are shown above each pair of boxplots. **d**. Hierarchical clustering on relative abundance of immune cell populations shows heterogeneity in sample composition within patients and controls. **e**. Scatter plot of proportion of monocyte and macrophage subsets and % of predicted forced vital capacity (FVC), Pearson correlation. Shaded area corresponds to the 95% confidence interval. **f**. Scatter plot of proportion of monocyte and macrophage subsets and the average fibrotic score on chest high-resolution computed tomography (HRCT), Pearson correlation. Shaded area corresponds to the 95% confidence interval.

We resolved six transcriptionally distinct TRAM clusters. TRAM-6 expressed genes encoding cytokines and chemokines (*IL1B*, *CCL3*, *CCL4*, *CCL20*, *CXCL8*, *CXCL10*, and *SOD2*). TRAM-5 expressed genes belonging to the metallothionein family (*MT1M*, *MT1G*, *MT1E*) (**Figure S1c**). We resolved three clusters of MoAM (MoAM-1, -2,-3). Macrophages in cluster MoAM-1 were the least mature and shared some genes with classical monocytes (*CCR2*, *CD300E*, *FCN1*, *CCL2*, *VCAN*). Macrophages in cluster MoAM-2 lacked expression of both *CCR2* and *CCL2* and included cells that expressed MoAM (*FCN1*, *VCAN*, *MSR1*) and TRAM (*FABP4*, *NUPR1*, and *INHBA*) markers. The transcriptional continuum of marker genes for these clusters suggested their progressive differentiation from classical monocytes into immature MoAM-1 into the more mature MoAM-2 as they adapt to long-term residence in the alveolar space (**Figure 1b**, **Figure S1c**). Cluster MoAM-3 was characterized by the expression of *SPP1*, *CHI3L1*, *PLA2G7*, *CHIT1*, and other genes associated with the development of pulmonary fibrosis and matched profibrotic macrophages reported in other fibrotic lung diseases^13,16,23,25^. Unlike alveolar macrophages, interstitial macrophages did not express genes indicating long-term adaptation to alveolar niche, such as *PPARG*, *FABP4*, and *MCEMP1* (**Figure 1b**, **Figure S1c**), suggesting that detection of interstitial macrophages in the alveolar compartment is a non-specific spillover from the interstitial compartment. We identified two clusters of classical monocytes characterized by expression of *FCN1*, *S100A12*, *CD300E*, *VCAN*, *CCR2* and lack of *MRC1*, *MSR1*, and *C1QA*. We labeled one cluster “Activated Monocytes” based on increased expression of cytokines and chemokines, including *CXCL10*, *CXCL11*, and *CCL8*.

We then compared the abundance of these cell populations between patients and controls. Profibrotic MoAM-3, MoAM-1, interstitial macrophages, monocytes, proliferating macrophages, *CCR7+* DC, and DC2 were significantly more abundant in patients compared to controls (**Figure 1c**). Of note, *SPP1+* MoAM-3 were virtually absent in controls. Hierarchical clustering by cell type abundance demonstrated substantial patient heterogeneity driven by differences in TRAM and MoAM subset proportions (**Figure 1d**). The abundance of MoAM-1, *SPP1+* MoAM-3, and interstitial macrophages negatively correlated with FVC, and positively correlated with fibrotic score on chest high-resolution computed tomography (HRCT) (**Figure 1e,f**). In contrast, an abundance of TRAM-2 and *MT1G+* TRAM-5 negatively correlated with fibrotic score on HRCT. No populations correlated significantly with the ground glass score on HRCT. Together, the abundance of profibrotic MoAM-3 detected by scRNA-seq was positively associated with functional and radiographic severity of SSc-ILD.

### Macrophage and T cell gene programs associate with SSc-ILD

We identified differentially expressed genes (DEGs) for each cell type between patients and controls. We found 5–40% of detected cell type-specific transcriptomes were differentially expressed in patients compared to controls (**Figure 2a,b**; **Table S2**). To interpret these gene expression changes, we performed gene set enrichment analysis (GSEA) using 50 hallmark gene sets from MSigDB (**Figure 2c**; **Table S3**). GSEA demonstrated significant enrichment for genes involved in “Oxidative Phosphorylation” in MoAM-1, MoAM-2, and T cells, excluding regulatory T cells (**Figure 2c**). The leading-edge genes for this gene set included genes encoding components of mitochondrial complex I, III and IV (*NDUFC2*, *COX4I1*, *COX5A*, *COX6A1*, *COX6B1*, *COX6C*, *NDUFA4*) and mitochondrial solute membrane transporters (*SLC25A5*, *SLC25A3*) (**Table S3**).

**Figure 2:**
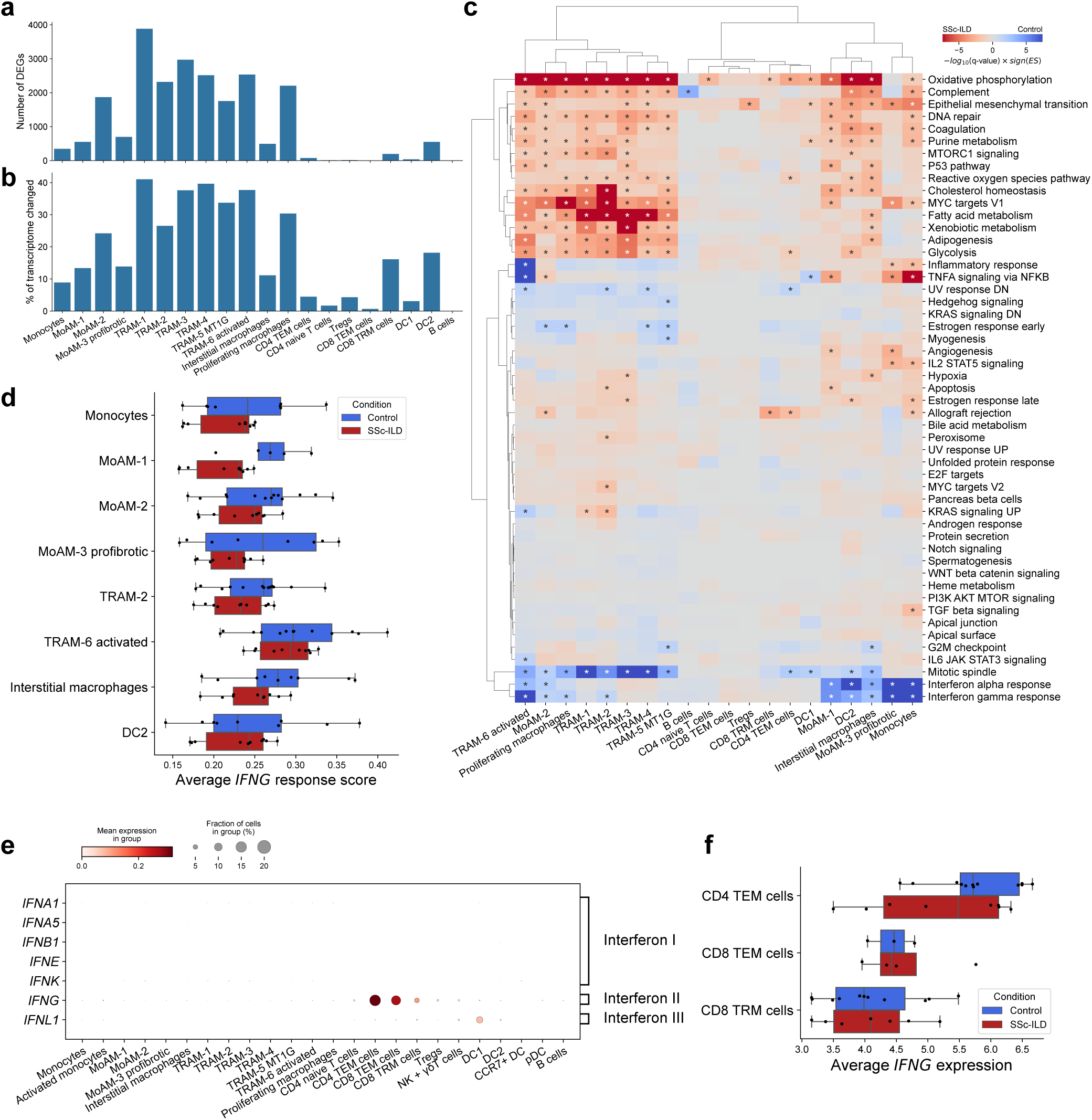
Macrophage and T cell gene programs associate with SSc-ILD. **a**. Number of differentially expressed genes (DEGs, *q*-value < 0.05) in different cell types between controls and patients. **b**. Percent of transcriptome within different cell types represented by DEGs. **c**. Heatmap illustrating enrichment for 50 hallmark gene sets from MSigDB in gene lists between controls and patients. Significant enrichments (*q*-value < 0.05) are indicated by asterisks. **d**. Average gene set score for genes from “interferon-gamma response” gene set in monocytes, macrophages, and DC2. *q*-value < 0.05 are shown next to pairs of boxplots (Mann-Whitney U tests with FDR correction). **e**. Dot plot illustrating expression of genes encoding type I, II, and III interferons in immune cells isolated from BALF. **f**. Expression of *IFNG* in T cell subsets isolated from BALF from controls and patients. *q*-value < 0.05 are shown next to pairs of boxplots (Mann-Whitney U tests with FDR correction).

Genes implicated in “Allograft Rejection” were significantly enriched in monocytes, MoAM-2, CD4 T effector memory (TEM) cells, and CD8 T resident memory (TRM) cells from patients. In monocytes, these included genes important for monocyte recruitment and maintenance in the niche (*CCL2*, *CCR2*, and *CSF1*), interferon response (*IFNGR2*), cell activation and signaling (*IL4R*, *CD40*, *CD86*, *IGSF6*, *LY86*, *TLR2*), matrix remodeling (*TIMP1*), and Fc receptor-mediated signaling (*FCGR2B*, *FGR*). In MoAM-2 these included *CCL2*, *CCR2*, *ICAM1* and *TGFB1*. In T cells, these included genes involved in cytokine signaling (*CCL4*, *CCL5*, *CXCR3*, *LTB*) and T cell maturation and function (*LCK*, *CD3D*, *CD3E*).

In both MoAM and TRAM subsets from patients compared to controls, gene sets “coagulation”, “complement”, “MYC targets”, “cholesterol homeostasis”, “epithelial mesenchymal transition” were significantly enriched. In all subsets of monocytes and MoAM, the gene set “TNFa signaling via NFKB” was enriched in patients compared to controls. Several gene sets were enriched only in specific cell types. For example, gene sets “fatty acid metabolism”, “glycolysis”, and “xenobiotic metabolism” were significantly enriched in all six TRAM subsets and in MoAM-2, while MoAM-1 subset in patients was enriched for “apoptosis” pathway.

A heightened interferon-response signature in the circulation is a hallmark of SSc and has been associated with improved response to therapy with myelosuppressive agents^26,27^. Surprisingly, in patients compared to controls, we found expression of “interferon-alpha response” and “interferon-gamma response” gene sets were reduced in DC2, monocytes, all MoAM subsets, and TRAM-2 and TRAM-6 subsets (**Figure 2c,d**). We then asked which immune cells in BAL fluid expressed interferons. Only DC1 had detectable expression of type III interferon *IFNL1*, and no cell types expressed detectable levels of type I interferon genes. Plasmacytoid dendritic cells (pDC) are a major source of type I interferon and have been suggested to play a role in the pathogenesis of SSc-ILD^28^. However, we did not detect any type I interferon expression in pDC in patients or controls (**Figure 2e**). CD4 TEM cells, CD8 TEM, and TRM cells expressed type II interferon *IFNG* (**Figure 2e**). Despite downregulation of the “interferon-gamma response” gene set in patients, *IFNG* expression in T cell subsets was similar between patients and controls (**Figure 2f**).

While TRAMs exist in multiple transcriptional states, we reasoned that they all should reflect disease-associated changes in their local microenvironment. Therefore, we focused on DEGs shared between all six TRAM subsets (704 upregulated and 349 downregulated genes in patients versus controls, **Figure S2a**). We found 18 upregulated gene ontology (GO) biological processes for upregulated genes, and none for downregulated genes (**Table S4**). The enriched GO processes included those involved in mitochondrial function and cellular respiration (**Figure S2b**). Similarly, GO processes related to mitochondrial function were identified in DEGs upregulated in patients for MoAM-1 (363 genes) and MoAM-2 (971 genes) (**Figure S2c**, **Table S5**).

We identified 310 upregulated and 182 downregulated DEGs in interstitial macrophages from patients (**Figure S2d**, **Table S2**). Unlike differential expression analysis in alveolar macrophage subsets, in interstitial macrophages GSEA identified “hypoxia” gene set enriched in patients and its leading edge included genes related to glycolysis (*ENO1*, *GPI*, *ALDOA*), glucose uptake (*SLC2A3*, also known as GLUT3), inflammation, phagocytosis, and tissue remodeling (**Table S3**). Expression of *CCL13*, which encodes a cytokine that recruits monocytes, eosinophils, basophils, and T cells via CCR2, CCR3, and CCR5, correspondingly, and *CCL18*, a cytokine that recruits T cells via CCR8 was upregulated in interstitial macrophages from patients (**Figure S2d**).

PCR-based biomarkers that can be applied to BAL fluid are already in clinical practice. Thus, we averaged RNA expression for all BAL cells in a given subject and determined the correlation between individual gene expression levels and pulmonary function test results. This analysis identified 42 genes that were significantly associated with pulmonary function (*q*-value < 0.01). Genes expressed in TRAM were correlated with a higher FVC, including *STIM1*, *ITGAL*. Genes expressed in *SPP1+* MoAM-3 or interstitial macrophages were correlated with low FVC, including *MARCKS*, *LGMN*, *FCGR2B*, *MAFB*, *EMP1* (**Figure S2e**, **Table S6**). These results suggested that detection of alveolar macrophage subsets using flow cytometry or PCR on bulk cellular RNA could be developed as a biomarker for SSc-ILD severity or response to therapy.

### Mycophenolate therapy is associated with transcriptomic changes in immune cells

As previously mentioned, four out of nine patients were on active mycophenolate therapy at the time of BAL. Mycophenolate inhibits inosine-5′-monophosphate dehydrogenase which is necessary for guanosine production in all cells. Resultant guanosine nucleotide depletion is thought to reduce the proliferation of T and B cells, and most studies attribute its anti-inflammatory effects to these changes^4,29^. The effects of mycophenolate on other immune cells, including monocytes and macrophages, are understudied^30^. Therefore, we performed differential gene expression analysis in lung immune cells between patients currently treated or untreated with mycophenolate. We detected 150–700 DEGs in TRAMs, MoAMs, DCs and the CD4 TEM subset, which comprised 5–11% of the total transcriptome in a specific cell type (**Figure S3a,b**, **Table S7**).

GSEA identified gene sets differentially enriched across cell types between patients prescribed mycophenolate in comparison to untreated patients (**Figure 3a**, **Table S8**). In TRAM-6 the absence of mycophenolate treatment was associated with “inflammatory response”, “interferon gamma response” and “TNFa signaling via NFkB”, with *CCL2*, *CXCL11*, *C3AR1*, *CXCL11*, *IRF8*, *ICAM1*, *NFKBIE* and *CCL20* among the leading edge genes (**Figure 3a**, **Table S8**). In profibrotic MoAM-3 the absence of mycophenolate treatment was associated with “allograft rejection” with *ITGAL*, *CCR5*, *ICAM1*, and *MMP9* among the leading edge genes (**Figure 3a**, **Table S8**). Several other gene sets related to metabolism and cell cycle were differentially enriched in monocytes and MoAM-2 from patients prescribed mycophenolate compared to those without. We detected no significant differences between lung immune cell type proportions in BAL fluid between patients prescribed mycophenolate therapy and not (**Figure S3c**).

**Figure 3:**
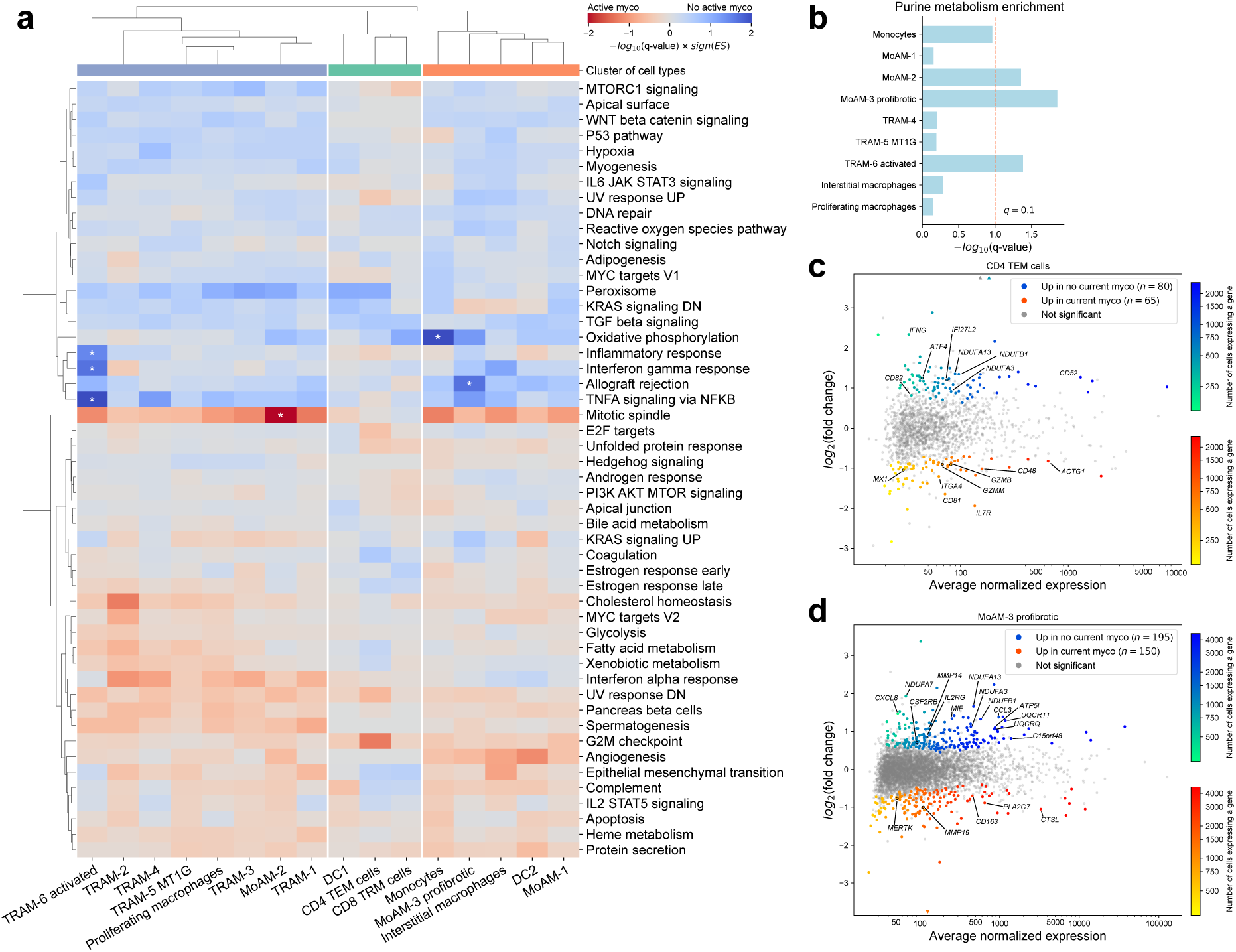
Mycophenolate therapy is associated with transcriptomic changes in immune cells. **a**. Heatmap illustrating enrichment for 50 hallmark gene sets from MSigDB in gene lists between patients on active therapy with mycophenolate and not. Significant enrichments (*q*-value < 0.05) are indicated by asterisks. **b**. Barplot of GO enrichment for “purine metabolism” gene set from KEGG database in DEGs upregulated with active mycophenolate therapy in macrophage subsets. **c**. MA plot of differential expression analysis between patients on active therapy with mycophenolate and not in CD4 TEM cells. Significant genes (*q*-value < 0.05) are highlighted in color. Selected genes of interest are labeled. Triangles indicate points that fall outside of the plotting window. **d**. MA plot of differential expression analysis between patients on active therapy with mycophenolate and not in profibrotic MoAM-3 cells. Significant genes (*q*-value < 0.05) are highlighted in color. Selected genes of interest are labeled. Triangles indicate points that fall outside of the plotting window.

We found the “Purine metabolism” gene set from KEGG database was significantly enriched in DEGs upregulated in patients prescribed mycophenolate in TRAM-6, MoAM-2, and MoAM-3 profibrotic clusters (**Figure 3b**). CD4 TEM cells had the highest IFNG expression, however, we did not detect significant enrichment for any gene sets in DEGs between patients prescribed mycophenolate and not. Therefore, we manually reviewed DEGs in this cell type. In patients prescribed mycophenolate, we detected significant downregulation of IFNG expression, as well as downregulation of genes involved in interferon signaling, mitochondrial respiration and immune cell regulation (*IFI27L2*, *ATF4*, *CD52*, *NDUFA3*, *NDUFA13*, *NDUFB1*, *CD82*). Conversely, CD4 TEM cells from patients prescribed mycophenolate demonstrated increased expression of genes involved in control of T cell survival, activation, and effector function: *IL7R*, *CD81*, *ACTG1*, *ITGA4*, *CD48*, *MX1*, *GZMB*, *GZMM* (**Figure 3c**, **Table S7**).

We also examined changes associated with mycophenolate therapy in profibrotic MoAM-3 and found downregulation of genes related to mitochondrial electron transport chain (*NDUFA7*, *NDUFA13*, *NDUFB1*, *NDUFA3*, *UQCR11*, *UQCRQ*, *C15orf48*), cytokine signaling (*CXCL8*, *CCL3*, *MIF*, *IL2RG*, *CSF2RB*), among others. In contrast, we found upregulation of genes related to heme metabolism, phagocytosis, phospholipid catabolism, including *CD163*, *MERTK*, *PLA2G7*, *MMP19*, and *CTSL* (**Figure 3d**, **Table S7**). Overall, these results suggest the effects of mycophenolate extend beyond its effects on T and B cell proliferation.

### Profibrotic *SPP1+* MoAM represent a pathologic monocyte-to-macrophage differentiation trajectory in SSc-ILD

In wild-type and humanized mice, classical monocytes that transmigrate from the circulation into the alveolar space rapidly differentiate into progressively more mature MoAM subsets and then develop a stable TRAM phenotype in response to signals from a healthy alveolar niche^31,32^. Consistent with these findings, we detected a small number of monocytes, MoAM-1, and MoAM-2 in BAL fluid from controls. Monocytes and MoAM-1 were more abundant in patients, suggesting increased recruitment (**Figure 1c**), while the abundance of the more mature MoAM-2 population was similar between patients and controls. In contrast, profibrotic *SPP1+* MoAM-3 were nearly exclusively found in patients (**Figure 1c**). This prompted us to ask if the differentiation trajectory toward *SPP1+* MoAM-3 was distinct from the typical monocyte-to-macrophage differentiation observed in the normal lung.

We used CellRank 2 to estimate cell transitions using pseudotime^33^. CellRank 2 identified seven stable terminal macrostates from the cell-cell transition matrix, which contained cells from both patients and controls (**Figure 4a**, **Figure S4a,b**). Since our analysis focused on monocyte to MoAM to TRAM differentiation rather than on transitions within TRAMs, we grouped TRAM-1, TRAM-2, TRAM-4 and TRAM-5 terminal states into a single TRAM terminal state and excluded interstitial macrophages. Next, we compared the average probability of differentiation into each terminal state for monocytes, MoAM-1, and MoAM-2 for cells from patients vs. controls. The probability of monocytes and MoAM-2 differentiation towards the combined TRAM terminal state was higher in samples from controls than samples from patients (**Figure 4b**). Conversely, the probability of monocytes and MoAM-2 differentiation into a profibrotic MoAM-3 terminal state was higher for patients versus controls (**Figure 4c**).

**Figure 4:**
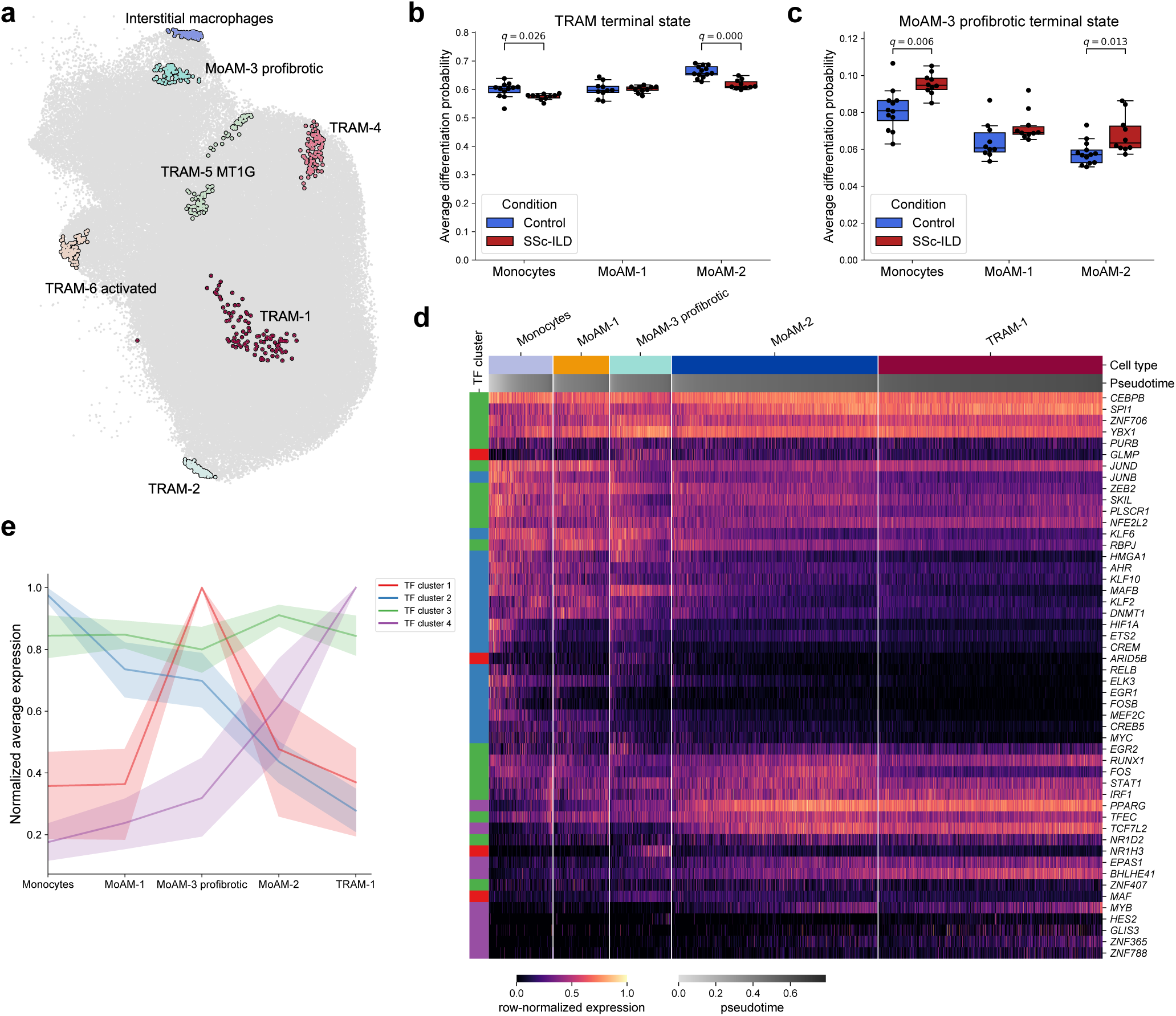
Profibrotic *SPP1+* MoAM represent a pathologic monocyte-to-macrophage differentiation trajectory in SSc-ILD. **a**. UMAP of macrophages and monocytes from **Figure 1a** with projected terminal states identified by CellRank 2. **b**. Average differentiation probability of monocytes, MoAM-1, and MoAM-2 subsets towards the combined TRAM terminal state. *q*-value < 0.05 are shown next to pairs of boxplots (Mann-Whitney U tests with FDR correction). **c**. Average differentiation probability of monocytes, MoAM-1, and MoAM-2 subsets towards the MoAM-3 profibrotic terminal state. *q*-value < 0.05 are shown next to pairs of boxplots (Mann-Whitney U tests with FDR correction). **d**. Heatmap of normalized expression of transcription factors in monocytes, MoAM subsets, and 50% of TRAM-1 selected randomly. Cells are ordered by pseudotime within each cluster, and expression is averaged over a window of 10 cells. **e**. Normalized average expression of transcription factor clusters for the cells in panel d. Shaded area represents 95% confidence interval.

Analysis of expression of transcription factors associated with monocyte-to-alveolar macrophage differentiation identified four patterns. While some transcription factors, including *CEBPB*, *SPI1*, and *RUNX1* (cluster 3) demonstrated stable expression across monocytes and macrophage subsets, other transcription factors demonstrated a continuous gradient of changes for monocyte to MoAM-1 to MoAM-2 to TRAM, for example downregulation of *MAFB*, *DNMT1*, and *HIF1A* (cluster 2) and upregulation of *PPARG* and *BHLHE41* (cluster 4). In contrast, transcription factors expressed in profibrotic *SPP1+* MoAM-3, such as *ARID5B*, *GLMP*, *MAF*, *NR1H3* (cluster 1) did not fit this trajectory (**Figure 4d,e**).

Ability to self-renew is one of the hallmarks of long-term adaptation to tissue residency. Since abundance of proliferating macrophages was increased in patients (**Figure 1c**), we analyzed the composition of this cluster and separated cells into TRAM- or MoAM-like. While the abundance of proliferating TRAM-like cells was not different between patients and controls, the abundance of MoAM-like cells was higher in patients (**Figure S4c**). Combined, these analyses suggest that MoAM-3 represent a distinct pathologic differentiation trajectory in patients with SSc-ILD and emphasize the importance of their abundance and gene signatures as biomarkers.

### TRAMs and profibrotic MoAMs occupy the same spatial niche in patients with SSc-ILD

To determine the location of profibrotic MoAM-3 in the fibrotic lung we performed single-cell spatial transcriptomic profiling of explanted lungs from four patients with end-stage SSc-ILD (five sections) and three lung donors (six sections) with two technical replicates for each sample (see **Table 3** for demographic and clinical characteristics of SSc-ILD and donor lungs). As expected, lungs from patients with SSc-ILD showed a cellular or fibrotic nonspecific interstitial pneumonia (NSIP) pattern, with loss of normal alveolar structure, bronchiolization, microscopic honeycombing, regions of fibrosis and inflammation, pleural thickening, thickened and hypertrophied blood vessel walls, and tertiary lymphoid structures (**Figure S5a**). Donor lungs were histologically normal (**Figure S5a**). Supervised iterative clustering on gene expression profiles resolved twelve epithelial, twelve stromal, ten endothelial, and 30 immune cell types, representing cell types and states previously reported in donor lungs and lungs from patients with pulmonary fibrosis (**Figure 5a**, **Figure S5b**, **Table S9**). Within macrophages, we resolved clusters that matched TRAM (*MARCO+SPP1–FOLR2–*), MoAM (*MARCO+SPP1+FOLR2–*), and interstitial macrophages (*MARCO–SPP1–FOLR2+*) (**Figure 5b**).

**Figure 5:**
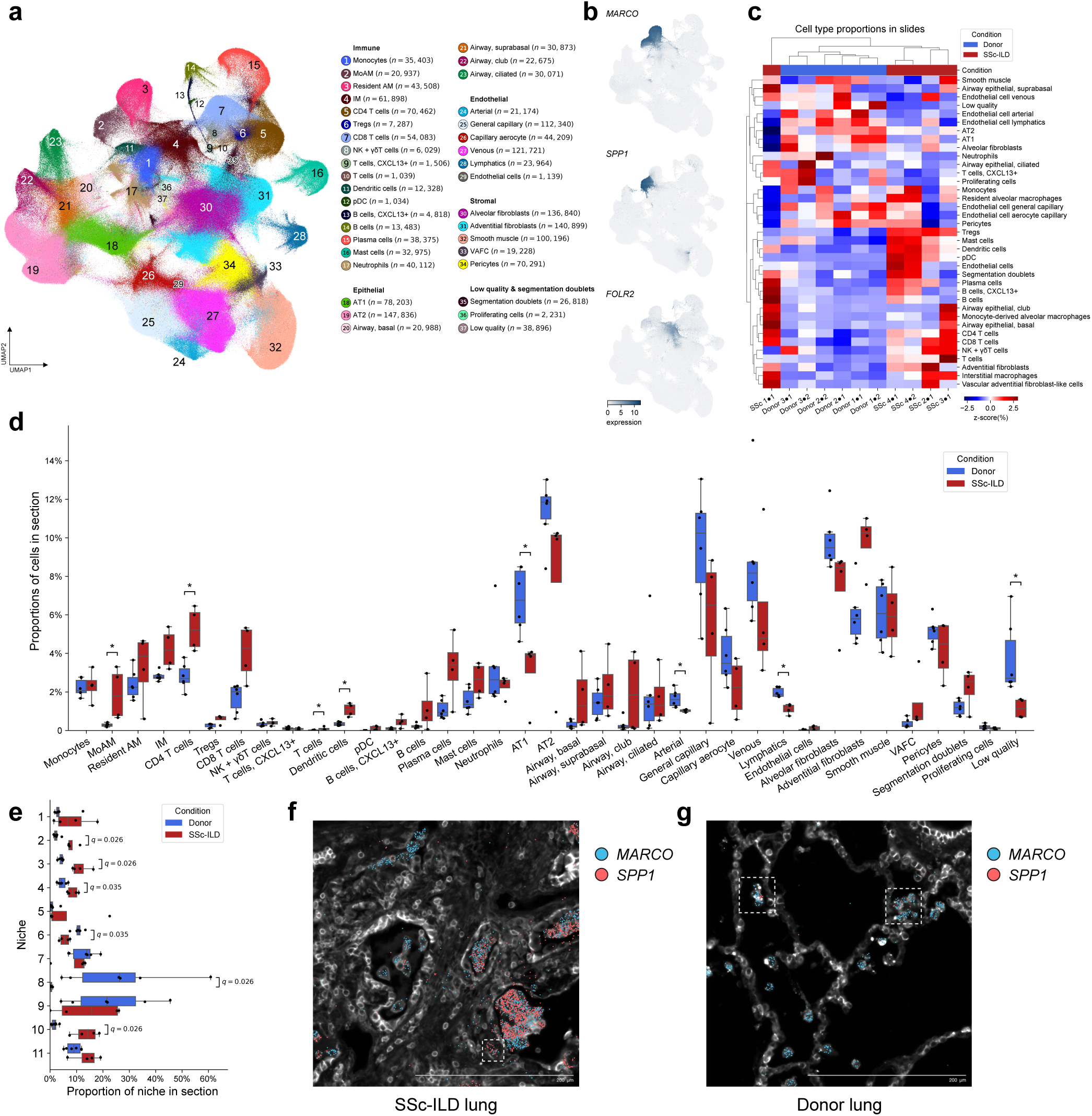
Single-cell spatial profiling identifies niche for profibrotic *SPP1+* MoAM. **a**. UMAP plot showing integrated analysis of single-cell spatial data from patients with SSc-ILD (*n* = 4) and donors (*n* = 3). **b**. Expression plot for markers of TRAM (*MARCO+SPP1–FOLR2–*), MoAM (*MARCO+SPP1+FOLR2–*), and interstitial macrophages (*MARCO– SPP1–FOLR2+*) on the UMAP from panel **a**. **c**. Hierarchical clustering of cell type proportions for 5 lung slides from patients with SSc-ILD and 6 donor lung slides. Each row is normalized. Ward linkage over Euclidean distances. **d**. Comparison of proportions of cell clusters shown in panel a between lungs from donors and patients with SSc-ILD. *q*-value < 0.05 are shown above each pair of boxplots, Mann-Whitney U tests with FDR correction. **e**. Comparison of proportions of spatial niches between lungs from donors and patients with SSc-ILD. *q*-value < 0.05 are shown above each pair of boxplots, Mann-Whitney U tests with FDR correction. **f**. Representative Xenium image showing localization of MoAM in both alveolar and interstitial compartments in the lung from patient with SSc-ILD. Xenium probes for marker genes shown as colored dots. **g**. Representative Xenium image showing localization of MoAM in alveolar epithelial niche in the donor lung. Xenium probes for marker genes shown as colored dots. Cell outlines are shown using boundary stain from Xenium cell segmentation kit.

**Table 3:**
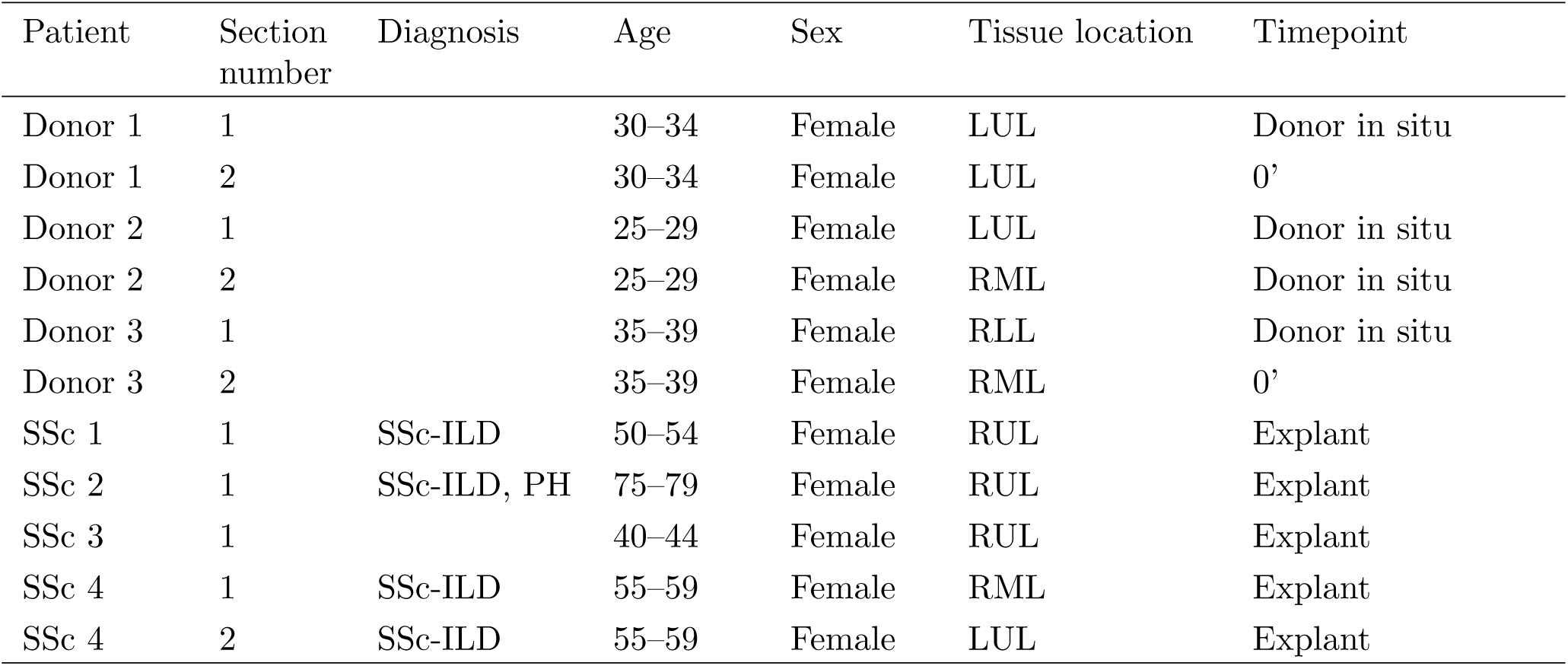
Tissue samples with patient demographics for single cell spatial transcriptomic analysis.

We observed high correlations in cell type compositions between technical replicates (**Figure S5c**). Hierarchical clustering of cell type proportions demonstrated that samples clustered based on disease status (SSc-ILD vs. donor) and that samples from patients with SSc-ILD were more heterogeneous (**Figure 5c**). Differential abundance analysis across major lineages demonstrated a decreased proportion of endothelial cells and expansion of immune cells in the lungs from patients (**Figure S5d**). Differential abundance analysis across clusters (**Figure 5a**) demonstrated a decreased proportion of alveolar epithelial type 1 (AT1) cells, endothelial arterial cells, and endothelial lymphatic cells in lungs from patients, while CD4 T cells, conventional DC2, and MoAM were more abundant in the lungs from patients (**Figure 5d**).

Niche analysis resolved eleven niches (**Figure S5e–j**; see **Table S10** for a description of the niche composition). Niche 6, representing connective tissue around vessels, and niche 8, representing parenchyma of alveolar space, were under-represented in the lungs from patients (**Figure 5e**). In contrast, niches 10 (intra- and extravascular immune cells), 4 (endothelium of lymphatics and blood vessels), 2 (alveolar macrophages in airspaces), and 3 (stromal cells) were expanded in patients (**Figure 5e**).

A fibrotic niche 3 was comprised of alveolar fibroblasts, adventitial fibroblasts, and smooth muscle cells. This niche had more diffuse distribution in the lungs from patients and was significantly more abundant in comparison to donor lungs where it was restricted to airways. A vascular niche 4 was represented by venous endothelial cells, lymphatic endothelial cells, and pericytes and was significantly expanded in the lungs from patients. An intra- and extravascular immune cells niche (niche 10) was enriched for T cells, B cells, plasma cells, and interstitial macrophages. Niche 10 was present around the airways in donor lungs but had more diffuse distribution in the lungs of patients.

We found most TRAM and MoAM in niche 2, which was restricted to alveolar and airway spaces. This niche was significantly expanded in the lungs from patients (**Figure 5e**). Moreover, we found that in the lungs from patients the majority (91.2%) of *SPP1+* MoAM were localized in the alveolar space (niche 2) and were intermixed with TRAM (**Figure 5f**, **Figure S5e,f**). The remaining MoAM were localized to the vascular immune cell niche (niche 11, 3.6%) or the intra- and extravascular immune cells niche (niche 10, 1%). Direct examination confirmed that at least some of these MoAM were localized to the interstitial space in patients (**Figure 5f**). In the donor lungs 39.2% of MoAM were found in the alveolar epithelial niche (niche 8), which included alveolar epithelial type 1 and type 2 cells, endothelial general capillary cells, aerocytes, and alveolar fibroblasts (**Figure S5e**), with the remaining 14.7% localized to niche 2, which was confirmed by direct examination of the images (**Figure 5g**). Thus, in the lungs of patients the vast majority of profibrotic MoAM are located in the alveolar space adjacent to the alveolar epithelium with only a few MoAM localized with stromal cells in the interstitial compartment.

### Transcriptionally distinct alveolar macrophage subsets co-localize within the same alveoli

We and others have reported transcriptional heterogeneity within alveolar macrophage subsets from healthy subjects and patients with lung fibrosis, including distinct clusters characterized by the expression of metallothionein genes (*MT1M*, *MT1G*, *MT1E*, *MT1H*, and others), or genes related to NF-kB signaling and the production of inflammatory cytokines (*TNFAIP6*, *CCL3*, *CCL4*, and others) (**Figure 1a**, **Figure S1c**)^16,22–24^. However, whether these transcriptionally distinct alveolar macrophage subsets populate specific alveoli or co-exist within the same alveolus is unknown. Therefore, we manually annotated 100–400 alveoli per donor lung section (**Figure S6a**) and analyzed the distribution of transcriptional subsets of alveolar macrophages. We detected on average 3.2 alveolar macrophages per single alveolus (**Figure S6b**). We found that alveolar macrophages expressing *CCL3* or *CCL4* were not localized to specific alveoli but were randomly dispersed across multiple alveoli and frequently co-localized with *CCL3–* and *CCL4–* alveolar macrophages (**Figure 6a,b**, **Figure S6c**). We observed the same pattern for *MT1H+* alveolar macrophages (**Figure S6c–e**). Although *CCL3* and *CCL4* encode chemokines MIP-1a and MIP-1b, respectively, which attract neutrophils, monocytes, and T cells via *CCR1*, *CCR4*, and *CCR5*, we did not detect accumulation of immune cells other than alveolar macrophages in alveoli harboring *CCL3+CCL4+* alveolar macrophages (**Figure 6a,c**). We found that the small number of MoAM in the donor lung were randomly distributed in the alveoli and did not have a distinct colocalization pattern with other immune cells (**Figure 6a,d**). In donor lung alveoli harboring alveolar type 1 (AT1) or alveolar type 2 (AT2) epithelial cells expressing CCL2— a major chemoattractant for classical monocytes—we did not detect significantly higher abundance of monocytes or MoAM (**Figure 6e**). Together, these findings suggest that ontogenetically and transcriptionally distinct alveolar macrophage subsets coexist within the same alveolar microniches in the healthy human lung rather and their detection in BAL fluid from healthy volunteers does not represent sampling from regions of microinjury.

**Figure 6:**
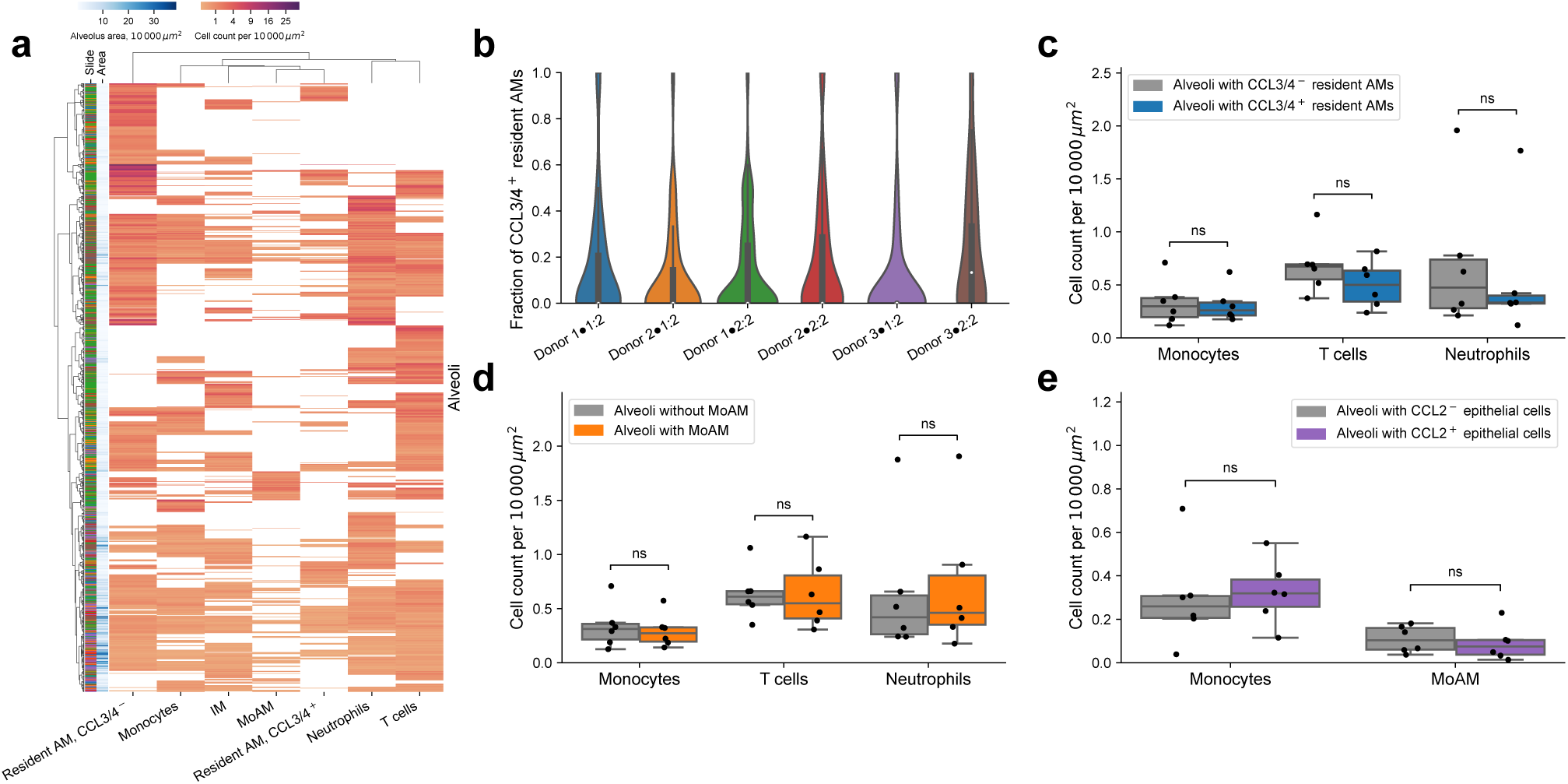
Transcriptionally distinct alveolar macrophage subsets are co-localized within the same alveoli. **a**. Hierarchical clustering on abundance of immune cell composition in the alveolar spaces in the donor lungs shows that *CCL3/4+* TRAM coexist together with *CCL3/4–* TRAM. Ward linkage over Euclidean distances. **b**. Violin plot of fractions of *CCL3/4+* TRAM out of all TRAM per alveolus across donor samples. White dot indicated median value, black bar corresponds to first to third quartile range. **c**. Comparison of the number of monocytes, T cells, and neutrophils in alveoli with and without *CCL3/4+* TRAM in the donor lungs. Pairwise Wilcoxon signed-rank tests with FDR correction. **d**. Comparison of the number of monocytes, T cells, and neutrophils in alveoli with and without MoAM in the donor lungs. Pairwise Wilcoxon signed-rank tests with FDR correction. **e**. Comparison of the number of monocytes and MoAM in alveoli with and without *CCL2* expression in AT1 or AT2 cells. Pairwise Wilcoxon signed-rank tests with FDR correction.

### Interstitial macrophages “spill” into alveolar space

Careful stereology studies showed that interstitial macrophages are at least as abundant as alveolar macrophages in non-diseased human lungs, but their role in lung disease is not well-studied. In naïve mice, investigators have identified at least two ontologically distinct subsets of interstitial macrophages: interstitial macrophages in the perivascular space that are long-lived and self-renewing and interstitial macrophages in the connective tissue and airways that are slowly replaced by circulating monocytes^34,35^. Perivascular macrophages are identified by expression of the hyaluronan receptor *Lyve1*. However, their human counterparts and their spatial localization are unknown. Therefore, we leveraged our spatial dataset to investigate the composition and localization of interstitial macrophages in donor lungs and lungs from patients.

Consistent with scRNA-seq data, interstitial macrophages were distinguished from monocytes and alveolar macrophages by relatively high expression of *STAB1*, *SELENOP* (*SEPP1*), and *FOLR2* (**Figure S5b**). In the donor lung, the majority of interstitial macrophages belonged to the vascular immune cell niche 11 (55.5%), the alveolar parenchyma niches 8 (22.0%) and 9 (7.9%), and walls of large vessels and pleura (niche 6, 5.3%) (**Figure S5e**).

We then compared the abundance and distribution of interstitial macrophages in patients and donors. We resolved eight transcriptional subsets of interstitial macrophages (IM-1-8, **Figure 7a,b**, **Figure S7a**). While all subsets of interstitial macrophages had detectable levels of *LYVE1* and *CSF1R*, cluster IM-1 had the highest expression of *LYVE1* (**Figure 7b**). Subsets IM-2-6 had similar expression of the core interstitial macrophage genes (*STAB1*, *FOLR2*, *CSF1R*) and differed only by the presence or absence of expression of specific cytokine or chemokine genes, such as *CCL2*, *CCL3/CCL4*, *CCL18*, and *CXCL8* (**Figure 7b**). Lastly, subset IM-7 was characterized by lower expression of *FOLR2*, *LYVE1*, and *VSIG4*, and increased expression of *ITGAX*. In donor and patient lungs, subset IM-2 (*FOLR2+*) was the most abundant (**Figure 7c**). Overall, patients had more interstitial macrophages compared to donors (**Figure 5d**). Specifically, IM-2 (*FOLR2+*) and IM-3 (*FOLR2+CCL18+*) subsets were increased, and the IM-4 (*FOLR2+CCL2+*) subset was decreased in patients versus control lungs (**Figure 7c**).

**Figure 7:**
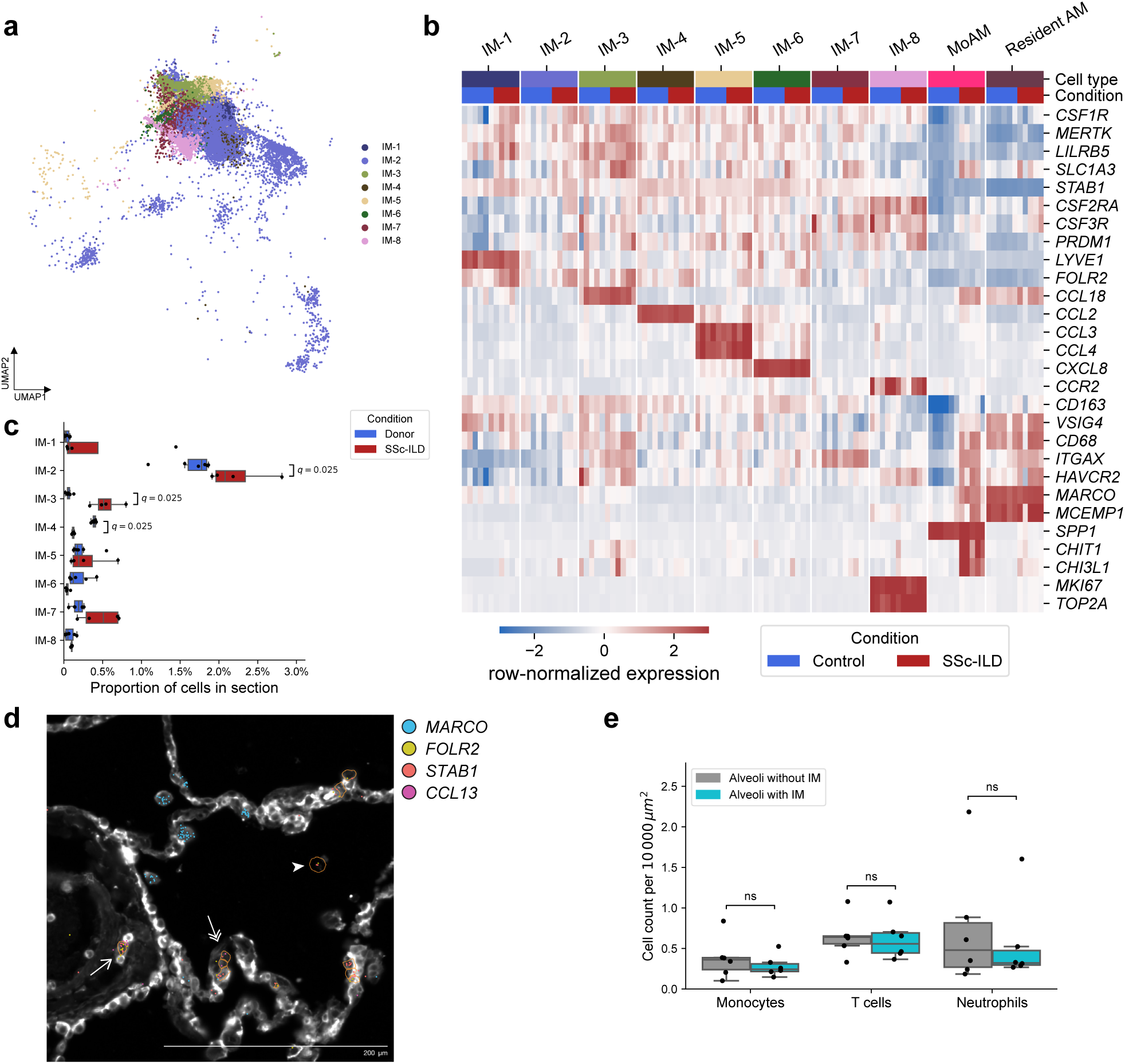
Interstitial macrophages “spill” into alveolar space. **a**. UMAP illustrating subsets of interstitial macrophages, reclustering of interstitial macrophage clusters from **Figure 5a**. **b**. Heatmap of expression of shared and unique genes among interstitial macrophage subsets and resident or monocyte-derived alveolar macrophages, each column represents a single donor or patient. **c**. Comparison of proportions of interstitial macrophage subsets between lungs from donors and patients with SSc-ILD. *q*-value < 0.05 are shown above each pair of boxplots, Mann-Whitney U tests with FDR correction. **d**. Representative Xenium image showing interstitial macrophages associated with blood vessel (arrow), alveolar parenchyma (double arrow), and alveolar space (arrowhead) of the donor lungs. Cell boundary stain from Xenium cell segmentation kit is shown. Each dot represents corresponding transcripts (see legend). Orange polygons represent interstitial macrophages as segmented by Xenium Ranger. **e**. Comparison of the number of monocytes, T cells, and neutrophils between alveoli with and without interstitial macrophages in the donor lungs. Pairwise Wilcoxon signed-rank tests with FDR correction.

BAL exclusively samples airspace and alveolar compartments. Thus, it is surprising that we and others identified interstitial macrophages in BAL fluid from patients and controls^22,23^. Spatial transcriptomics confirmed the presence of interstitial macrophages in airspaces in lungs from patients and donors, where they were sometimes associated with alveolar epithelium (**Figure 7d**). Detection of interstitial macrophages in the alveolar space of donor lungs was not associated with increased abundance of other immune cells (**Figure 6a**, **Figure 7e**). Together with the lack of expression of genes associated with long-term residency in the alveolar space (**Figure 1b**), these findings suggest that interstitial macrophages “spill” into the alveolar space in the normal and diseased lung where they can be sampled bronchoscopically.

### Single-cell genomics identifies therapeutic targets for SSc-ILD

Biological therapies including B cell-depleting therapies and tocilizumab, a monoclonal antibody blocking IL-6 receptor, have demonstrated positive signal in clinical trials in patients with SSc-ILD, and more biologics are currently in clinical trials^4^. Therefore, we compiled a list of biologic therapies currently approved for the treatment of SSc-ILD or undergoing testing in clinical trials and evaluated the expression of their targets in lung tissue. We excluded drugs that target B cell survival, TGF-β or its activating integrins, and broad-spectrum kinase inhibitors. We found 14 drugs in 16 trials on ClinicalTrials.gov that matched these criteria (**Table S11**). Two drugs targeted enzymes, five drugs targeted receptors, and 10 drugs targeted ligands. Next, we asked whether targets of these drugs had broad or cell-type-specific expression patterns, whether they were differentially expressed between cells isolated from the lungs of patients compared to controls in publicly available datasets from patients with end-stage SSc-ILD^12,15,36^ (**Figure S8a,b**, **Table S12**).

Out of 23 genes encoding targeted ligands, receptors, or enzymes, 11 had significant differences in expression between conditions in at least one cell type in the end-stage whole-tissue lung dataset (**Figure 8a**, **Figure S8c**, **Table S13**). Expression of *TNFSF13B*, which encodes a key B cell differentiation factor known as BAFF or BlyS (targeted by belimumab), was significantly upregulated in alveolar fibroblasts, CD8 T cells, plasma cells, and interstitial macrophages from patients. This finding suggests that fibrotic lung might serve as a source of factors that could support pathogenic B cells. In contrast, expression of genes encoding BAFF receptors (*TNFRSF13B*, *TNFRSF13C*, *TNFRSF17*) was not significantly different between controls and patients, including in B and plasma cells. Expression of *NRP2*, which encodes neuropilin-2 involved in myeloid cell differentiation and activation (targeted by efzofitimod), was significantly upregulated in secretory epithelial cells and profibrotic *SPP1+* MoAM-B from patients. Expression of *GUCY1A3*, one of the enzymes targeted by avenciguat, an oral soluble guanylate cyclase activator, was increased in adventitial and alveolar fibroblasts from patients, and decreased in venous endothelial cells. While expression of *TNFSF4*, which encodes OX40L and targeted by amlitelimab, was not different between patients and controls, expression of its receptor *TNFRSF4*, was downregulated in secretory airway epithelial cells from patients.

**Figure 8:**
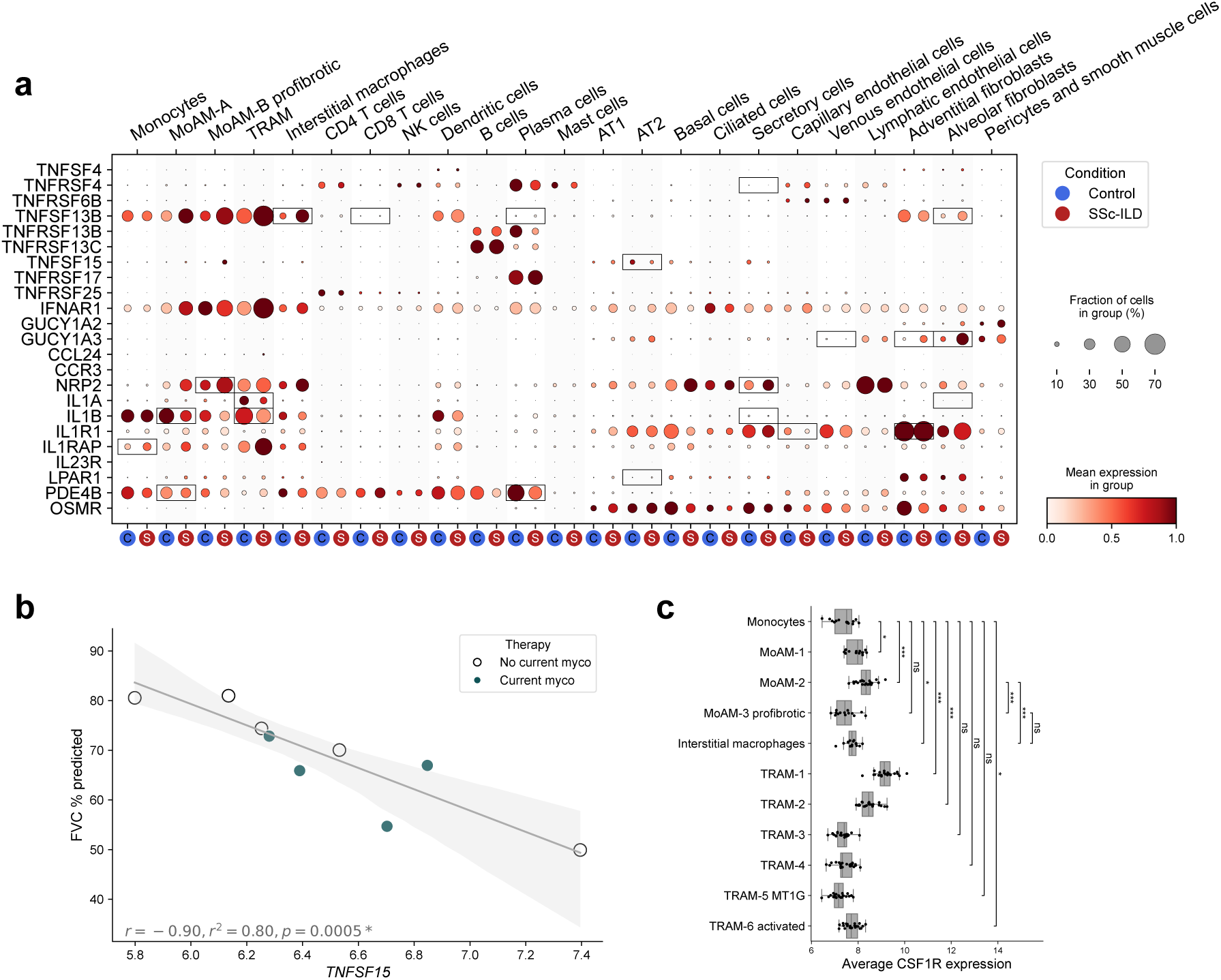
Pharmacological targets in patients with SSc-ILD. **a**. Dot plots illustrating expression of genes encoding ligands, receptors, or enzymes that are currently in clinical trials for patients with SSc-ILD. Reanalysis of publicly available data from patients with end-stage SSc-ILD. Significant differentially-expressed genes are outlined (see **Figure S8c,d**). **b**. Correlation between average expression of *TNFSF15* in BALF and % of predicted forced vital capacity (FVC). Pearson correlation. Shaded area corresponds to the 95% confidence interval. c. Expression of *CSF1R* in subsets of alveolar and interstitial macrophages from BALF. Mann-Whitney U tests with FDR correction, ns: not significant, * *q*-value < 0.05, ** *q*-value < 0.01, *** *q*-value < 0.001.

Expression of the gene encoding IL-1b (targeted by genakumab) was decreased in MoAM-A, TRAM, and secretory airway epithelial cells from patients. Similarly, expression of the gene encoding IL-1a was significantly decreased in alveolar fibroblasts and TRAM from patients. And while the expression of *IL1R1*, encoding interleukin-1 receptor type 1, was downregulated in capillary endothelial cells, it was significantly upregulated in adventitial fibroblasts from patients. Expression of *IL1RAP*, encoding interleukin-1 receptor accessory protein, which is important for the signaling of the interleukin-1 family of cytokines, was significantly upregulated in monocytes from patients.

LPAR1 signaling has been causally associated with pulmonary fibrosis, and an LPAR1 antagonist slowed the rate of FVC decline in a phase 2 clinical trial. While expression of *LPAR1* in AT2 cells was low, it was significantly upregulated in patients. In contrast, expression of *LPAR1* in stromal cells was not different between controls and patients. Nerandomilast, a PDE4B inhibitor, has also significantly slowed the rate of FVC decline in patients with idiopathic pulmonary fibrosis and non-IPF ILD^37,38^. Preclinical studies reported that PDE4B inhibitors act on both stromal and immune cells during pulmonary fibrosis. While expression of *PDE4B* was not different in stromal cells between controls and patients, we found that it was significantly downregulated in plasma cells and MoAM-A from patients.

The expression of *TNFSF15*, which encodes TL1A (targeted by tulisokibart, FG-M701, and several other monoclonal antibodies), was significantly downregulated in AT2 cells from patients. While expression of *TNFSF15* in immune cells was limited to macrophage subsets, it was not different between controls and patients with end-stage disease. In contrast, we found that in our BAL dataset, expression of *TNFSF15* in all cells captured in BAL fluid (pseudobulk) negatively correlated with FVC (**Figure 8b**).

Lastly, an antibody targeting the CSF1R, axatilimab, was reported to reverse established fibrosis in patients with refractory chronic graft versus host disease^11^. We found that expression of *CSF1R* increased as monocytes differentiated into MoAM-1 and MoAM-2 subsets (**Figure 8c**). However, the expression of *CSF1R* was lower in profibrotic MoAM-3 when compared to the MoAM-2 subset, and was not different from expression in monocytes or interstitial macrophages. While TRAM-1 and TRAM-2 subsets had increased expression of *CSF1R*, other TRAM subsets (TRAM-3-6) had *CSF1R* expression comparable to monocytes. In addition, TRAM are known to be critically dependent on CSF2R/CSF2 signaling^39^ and thus, maybe resistant to therapy with axatilimab.

## Discussion

We and others have causally linked the recruitment of profibrotic monocyte-derived alveolar macrophages to lung fibrosis development and maintenance in animal models^5,6,8,10^. The importance of monocyte-derived alveolar macrophages in human lung fibrosis remained uncertain until a recent phase 2 trial demonstrated that axatilimab, a monoclonal antibody targeting the CSF1 receptor, reversed established lung fibrosis in patients with chronic graft-versus-host disease refractory to other therapies^11^. As scRNA-seq data from lung explants of patients with pulmonary fibrosis localize expression of *CSF1R* exclusively to monocytes, macrophages and dendritic cells in the normal and fibrotic human lung, these results causally implicate one or more of these cell populations in pulmonary fibrosis.

In this study, we sampled the alveolar space in patients with carefully phenotyped SSc-ILD and used scRNA-seq to quantify the abundance and assess the transcriptomic profile of immune cell populations in the alveolar space. Our analysis identified a population of profibrotic monocyte-derived alveolar macrophages that were virtually absent in healthy volunteers and were expanded in patients with SSc-ILD. The abundance of these profibrotic monocyte-derived alveolar macrophages was associated with the severity of lung fibrosis as measured by HRCT and PFTs. Trajectory analysis suggested these cells might represent a terminal differentiation pathway from classical monocytes, distinct from the normal differentiation of monocytes through non-fibrotic monocyte-derived alveolar macrophage intermediate into mature, tissue-resident alveolar macrophages. We further identified interstitial macrophages in the alveolar space that were expanded in patients with SSc-ILD relative to healthy controls and associated with disease severity.

We used spatial transcriptomics to localize profibrotic monocyte-derived alveolar macrophages within the fibrotic lung and to compare the localization of interstitial macrophages in the normal and fibrotic lung. In agreement with previous reports, we found that profibrotic monocyte-derived alveolar macrophages were nearly exclusively confined to the alveolar space, with only rare cells in the interstitium, in direct contact with fibroblasts. These results suggest that profibrotic monocyte-derived alveolar macrophages may interact with the alveolar epithelium to drive injury or slow repair. This would create a vicious cycle of epithelial injury and failed repair, that promotes the ongoing recruitment of profibrotic monocyte-derived alveolar macrophages^9^. Alternatively, profibrotic monocyte-derived alveolar macrophages may exert their fibrotic effects on fibroblasts in transit, as they migrate into alveolar space via the interstitium.

We identified a substantial population of resident alveolar macrophages (∼5% of all cells in BAL in healthy volunteers), described by us and others, that express genes encoding chemokines and cytokines^22–24^. Our spatial analysis did not detect accumulation of inflammatory cells or activation of inflammatory signaling pathways in alveoli from normal lungs harboring these cells. This finding suggests these transcriptional states of resident alveolar macrophages in healthy subjects may represent primed transcription with translational repression, a phenomenon that has been described for T cells^40,41^.

Lung regions that normally harbor interstitial macrophages, in particular the areas around the vasculature, were expanded in patients with SSc-ILD with a corresponding increase in the abundance of interstitial macrophages. Importantly, vascular injury and perivascular lymphocytic infiltrates are a hallmark of SSc^2^. The interstitial macrophages sampled by the BAL procedure did not express markers indicating adaptation to the alveolar space. This suggests that they are spilling into alveoli from the interstitium, where they are sampled by the BAL procedure. These interstitial macrophages had transcriptomic profiles distinct from profibrotic monocyte-derived alveolar macrophages, providing insights into pathological processes in the lung interstitium from BAL samples.

Our findings have translational implications. First, because the risk of BAL procedure is relatively low^42^, quantification of profibrotic monocyte-derived alveolar macrophages might be used to inform a flow cytometry or PCR-based biomarker to monitor the response to therapy. Indeed, we performed many of our research bronchoscopies concurrent with scheduled esophagogastroduodenoscopy, a common clinically indicated procedure to assess esophageal dysfunction in patients with SSc. Second, the transcriptome of both tissue-resident and monocyte-derived alveolar macrophages varies as a function of mycophenolate treatment, perhaps providing insights into mechanisms explaining its efficacy in patients with SSc-ILD. Expanding this analysis to other therapies might provide similar insights. Third, our data support the hypothesis that axatilimab exerts its antifibrotic effect in the lung by targeting profibrotic monocyte-derived alveolar macrophages or interstitial macrophages, supporting its evaluation in future clinical trials for SSc-ILD^43^. Fourth, our data can be used to localize ligand-receptor interactions that are the target of current and proposed therapies for patients with SSc-ILD. Finally, our data suggest vasculopathy—a well-recognized early hallmark of SSc^44^—accompanied by expansion of interstitial macrophages^45^ might secondarily affect the alveolar epithelium, linking it to other causes of lung fibrosis that are genetically linked to failed repair of the alveolar epithelium^46^.

Our study has several limitations. First, our data are observational and cannot be used to infer causality. The incorporation of BAL sampling into clinical trials of new or existing therapies could address this limitation. Second, our spatial analysis and analysis of expression of genes encoding targets of drugs currently in trials in patients with SSc-ILD were limited to patients with end-stage lung fibrosis. Early, possibly druggable and reversible events important to disease pathogenesis might have resolved at the time of sampling. Third, we rely exclusively on transcriptomic analysis for these studies. Future studies using multiplexed protein approaches (e.g., CITE-seq or spatial proteomics) can address these limitations. In summary, the results of this paper support the hypothesis that the abundance and gene expression profiles of profibrotic monocyte-derived alveolar macrophages in BAL fluid from patients with SSc-ILD may be a biomarker of ILD severity and potential therapeutic target.

## Methods

### Human subjects

All research involving human participants was approved by the Institutional Review Board of Northwestern University or the Yale University Human Investigation Committee. Healthy volunteers were enrolled in studies STU00206783 and STU00214826 at Northwestern University, Pro00088966 and Pro00100375 at Duke University. Patients with SSc-ILD were enrolled in studies STU00202557, STU00218939, STU00204897, STU00002669, and STU00207083 at Northwestern University and HIC2000024862 at Yale University. All study participants provided informed consent, including consent for data sharing. Clinical data were extracted from the electronic health record using SQL queries from Northwestern Medicine Enterprise Data Warehouse and manual chart review by study-associated physicians and clinical coordinators.

A total of 9 patients with SSc, both with and without radiographically confirmed ILD (67% female, median age 54 (interquartile range 38–60) years were enrolled after undergoing evaluation at North-western Memorial Hospital and Yale New Haven Hospital between September 2018 and January 2022. Bronchoscopy was performed as a research procedure. Healthy controls were taken to the bronchoscopy suite at Northwestern Memorial Hospital, Yale New Haven Hospital, or Duke University Hospital. Detailed demographics and clinical characteristics of the study population are presented in **Table 1** and **Figure S1a**.

For single-cell spatial analysis, we used explanted lung tissue from patients with end-stage SSc-ILD undergoing lung transplantation. Histologically normal donor lung tissue obtained during the lung graft resizing was used as a control. These patients were enrolled in the study STU00212120 at Northwestern University.

Diagnosis of SSc was made based on 2013 ACR/EULAR criteria^47^ and was confirmed by a study-associated rheumatologist (C.R.; M.E.H.). SSc-ILD diagnosis was adjudicated by review of radiology reports from chest CT by experts in ILD (A.J.E.; J.E.D.; B.C.B. or J.P. and M.G.) utilizing a three-reader method that has been previously described^48,49^. SSc-ILD diagnosis was confirmed by a thoracic radiologist (H.S. or A.N.R.) who reviewed the associated chest CT imaging. HRCT scores were derived by study-associated radiologists (H.S. and A.N.R.) using the methodology described by Kazerooni et al. Pulmonary function test results were converted to % predicted based on a race-neutral reference from 2022 (see code).

### Bronchoscopy and BAL

Bronchoscopic BAL was performed in patients and healthy volunteers in the bronchoscopy suite as previously described^23,50^. Patients or healthy volunteers were given sedation and topical anesthesia at the discretion of the physician performing the bronchoscopy. Lavage was directed to lobes with higher GGO and fibrosis scores; 90–120 ml of saline was instilled into the segment of interest and aspirated back, with the first 5 ml of return discarded. BALF collected at Northwestern was processed fresh within 2 hours after collection. BALF samples collected at Yale and Duke were filtered through a 70-μm cell strainer, and mixed with HypoThermosol solution (Biolife Solutions, cat #101104) at 1:1 ratio and shipped overnight for further processing on wet ice to North-western University. BALF samples were processed using either “no FACS” or “FACS” protocol.

### BALF sample processing for scRNA-seq using “no FACS” protocol

BALF samples were filtered through a 70-μm cell strainer, pelleted by centrifugation at 400 rcf for 10 min at 4°C, followed by hypotonic erythrocyte lysis using 1 ml of PharmLyse solution (BD) for 2 min, followed by another wash with 10 ml of 0.5% BSA in PBS and centrifugation. Cell pellet was resuspended in 0.5% BSA in PBS at the final concentration 1000 cells/ul. Cell concentration and viability were determined using K2 Cellometer (Nexcelom) with AO/PI reagent and cells were loaded on 10x Genomics Chip A with Chromium Single Cell 3′ V2 gel beads and reagents (10x Genomics) aiming to capture 5,000–7,000 cells per library. Libraries were prepared according to the manufacturer’s protocol (10x Genomics, CG000052_RevB).

### BALF sample processing for scRNA-seq using “FACS” protocol

BAL fluid samples were filtered through a 70-μm cell strainer, pelleted by centrifugation at 400 relative centrifugal force (rcf) for 10 min at 4 °C, followed by hypotonic lysis of red blood cells with 2 ml of PharmLyse (BD Biosciences) reagent for 2 min. Lysis was stopped by adding 13 ml of MACS buffer (Miltenyi Biotech). Cells were pelleted again and resuspended in 100 μl of a 1:10 dilution of Human TruStain FcX (BioLegend) in MACS buffer and a 10-μl aliquot was taken for counting using a K2 Cellometer (Nexcelom) with Acridine Orange (AO)/Propidium Iodide (PI) reagent. The cell suspension volume was adjusted so the concentration of cells was always < 5 × 107 cells ml−1 and the fluorophore-conjugated antibody cocktail was added in a 1:1 ratio. The following antibodies were used (antigen, clone, fluorochrome, manufacturer, catalog no., final dilution): CD4, RPA-T4, BUV395, BD, 564724, 1:40; CD19, HIB19, BUV395, BD, 740287, 1:40; CD25, 2A3, BUV737, BD, 564385, 1:20; CD56, NCAM16.2, BUV737, BD, 612766, 1:20; HLA-DR, L243, eFluor450, Thermo Fisher Scientific, 48-9952-42, 1:40; CD45, HI30, BV510, BioLegend, 304036, 1:20; CD15, HI98, BV786, BD, 563838, 1:20; CD3, SK7, PE, Thermo Fisher Scientific, 12-0036-42, 1:20; CD127, HIL-7R, PECF594, BD, 562397, 1:20; CD206, 19.2, PECy7, Thermo Fisher Scientific, 25-2069-42, 1:40; CD8, SK1, APC, BioLegend, 344721, 1:40; CD14, M5E2, APC, BioLegend, 301808, 1:40; and EpCAM, 9C4, APC, BioLegend, 324208, 1:40. After incubation at 4 °C for 30 min, cells were washed with 5 ml of MACS buffer, pelleted by centrifugation and resuspended in 500 μl of MACS buffer with 2 μl of SYTOX Green viability dye (Thermo Fisher Scientific). Cells were sorted on a FACS Aria III SORP instrument using a 100-μm nozzle. Cells were sorted into 300 μl of 2% bovine serum albumin (BSA) in Dulbecco’s phosphate-buffered saline (DPBS) and immediately after sorting pelleted by centrifugation at 400 rcf for 5 min at 4 °C, resuspended in 0.5 BSA in DPBS to 1,000 cells μl−1 concentration. Concentration was confirmed using a K2 Cellometer (Nexcelom) with AO/PI reagent using the ‘Immune cells low RBC’ program with default settings and cells were immediately used for scRNA-seq. Cells were loaded on 10x Genomics Chip A with Chromium Single Cell 3′ V2 gel beads and reagents or 10x Genomics Chip B with Chromium Single Cell 3′ V3 gel beads and reagents (10x Genomics) aiming to capture 5,000–7,000 cells per library. Libraries were prepared according to the manufacturer’s protocol (10x Genomics, CG000052_RevB or CG000183_RevB).

Analysis of the flow cytometry data was performed using FlowJo v.10.7.1. using a sequential gating strategy reported in our previous publications and jointly reviewed by three investigators (Z.Y., S.S. and A.V.M.). A fraction of cells that was not definitively resolved by our panel was labeled ‘others’. Relative cell-type abundance was calculated as a percentage of all singlets/live/CD45+ cells.

### Statistical methods

No statistical method was used to predetermine sample size. The experiments were not randomized. The Investigators were not blinded to allocation during experiments and outcome assessment. All statistics in the manuscript are reported as specified in the figure legends. When multiple hypothesis tests were performed, the false discovery rate (FDR) was controlled using the procedure of Benjamini and Hochberg. A significance level of 0.05 was used for all tests, unless indicated otherwise.

### Figure and manuscript preparation

Plotting was performed in python using matplotlib and seaborn, for specific details see below and code. Figures were assembled in Adobe Illustrator. Spatial transcriptomics representative images were taken with Xenium Explorer software. Images of H&E sections for spatial transcriptomics were taken with NDP.view2 software from Hamamatsu. Manuscript was assembled with Typst.

### Single-cell RNA-seq analysis

Data were processed using Cell Ranger 3.1.0 (10x Genomics), reads were mapped to GRCh38.84 reference genome (10x Genomics Cell Ranger Human 1.2.0 GRCh38 reference). Data were processed using Scanpy 1.7.2^51^, and multisample integration was performed with scvi-tools 0.14.0^52^. The scVI models was constructed on 1000 HVGs with the hyperparameters `n_layers = 2, dropout_rate = 0.2`, and `n_latent = 10`, and were trained using the settings `max_epochs = 400, check_val_every_n_epoch = 2`, and `early_stopping = True`. Default hyperparameters and settings were used otherwise. An initial round of Leiden clustering using the function `sc.tl.leiden` was performed on the integrated BAL object with a resolution of 0.75. Clusters characterized by low number of detected genes and transcripts and high percentage of mitochondrial genes were removed. Clusters containing doublets were identified as clusters simultaneously expressing lineage-specific marker genes (for example, *C1QA* for macrophages and *CD3G* for T cells) and excluded. Cell types were identified by marker genes, computed using the `sc.tl.rank_genes_groups` function with the settings `method = “t-test”, n_genes = 200`, and default settings otherwise. For more details, see code.

Differential abundance analysis was performed on fractions of identified cell clusters in each BAL sample out of the whole sample, and the comparison was done with Mann-Whitney U tests. Correlation analysis with fibrosis scores and pulmonary function readouts were done with Pearson correlation.

### Differential expression analysis

To take advantage of the multiple subjects in each condition and avoid p-value inflation inherent to approaches where each cell is treated as an independent observation, we summed RNA transcript counts for each subject on per cell type level (pseudobulk approach). A sample was created if it contained at least 50 cells for a given cell type. Differential expression analysis was performed in R 4.1.1 using DESeq2 1.34.0^53^. A ‘local’ model of gene dispersion was used; default settings were used otherwise. Differentially expressed genes were those with q < 0.05 (Wald test with FDR correction). To help identify genes of interest, we applied two filtering criteria. First, we removed genes encoding ribosomal proteins, non protein-coding genes, genes mapping to NCBI accession IDs, and others (see code) from the list of differentially expressed genes. The second criterion was used to exclude genes lacking robust expression detection in our data. For each gene and cell type, we only kept genes that were expressed in at least 20 cells in at least 2 samples in both controls and patients with SSc-ILD groups, and which were expressed at the level of 50 average expression on DESeq2-reported baseMean scale. After removing the genes according to the two criteria above, we computed FDR-corrected p-values for the remaining genes, and report those.

To add interpretability to differential gene expression analyses results, we performed GSEA on the lists of filtered genes for each cell type, ordered by their log2FoldChange estimated by DESeq2. We used 50 hallmark gene sets from MSigDB, followed by FDR correction across all gene sets and all cell types (Figure 2c, Figure 3a). Separately, GO enrichment was performed on GO biological processes gene sets on significantly different genes, with FDR correction.

### Pseudotime estimation of cell transitions using CellRank 2

For differentiation trajectory analyses of macrophage populations in BALF we used Cellrank 2.0.4^33^. First, we selected only macrophage subsets from BALF single-cell object and estimated diffusion components with scanpy diffmap function. After manual inspection we identified diffusion component 2 as reflecting monocyte-to-macrophage differentiation, and picked a monocyte cell with maximum value for this component as the root node for pseudotime estimation. We used scanpy dpt function with default parameters to estimate pseudotime, and used PseudotimeKernel from cellrank to compute transition matrix for the cells. Next we used GPCCA estimator to compute Schur decomposition of the transition matrix with 40 components, and selected 7 macrostates after examining eigenvalues for the components. We manually combined TRAM terminal states since we were not interested in estimating transitions between resident macrophages, and computed cell fate probabilities using cellrank compute_fate_probabilities function with default parameters. Finally, we averaged cell fate probabilities on cell type and sample level, and compared them with t tests followed by FDR correction.

For transcription factor and differentiation trajectory analysis, we used the list of human transcription factors from Lambert et al.^54^. We selected macrophage subsets from BALF single-cell object. From the list of transcription factors we kept those that were expressed in at least 100 cells among macrophages in our data, and were either among marker genes for at least one cluster, or significantly differentially expressed in any cluster between samples from patients with SSc-ILD and controls. This gave us a list of 50 transcription factors. We then estimated pseudotime with scanpy diffmap function using the monocyte with the highest expression of *HIF1A* as the root. For Figure 4d we ordered cell clusters by average expression of known macrophage maturation transcription factor *PPARG*, ordered cells by pseudotime, and averaged expression for visualization using moving average over 10 cells.

To cluster transcription factors by their profile across macrophage subsets, we computed average normalized expression of each transcription factor for each cell cluster, and normalized the averages by maximum value. Then we used k-means clustering from scikit-learn package (with `n_init=1000` parameter) on 2 to 10 clusters and picked the best number of clusters (4) using average silhouette score.

### Single-cell spatial transcriptomics

Lung tissue was obtained at the time of transplant and stored on ice until processing. An approximately 10×10×10 mm piece of lung tissue was fixed in 10% neutral buffered formalin for 48 hours at 4 °C, followed by storage in 70% EtOH at room temperature, followed by series of ethanol and xylene dehydration steps and embedding in paraffin using Sakura Tissue-Tek VIP 5 instrument at 60 °C. Tissue blocks were rehydrated in an ice bath for 30 min and sectioned on Leica HistoCore Biocut microtome at 5 um thickness using the “Easy Does It” mode. After discarding the first 5 sections, the tissue sections were floated in a 42°C water bath, mounted onto Xenium slides, and baked at 42°C for 3 hours, followed by deparaffinization, decrosslinking, probe hybridization, ligation, and amplification using Xenium *In Situ* Gene Expression workflow (CG000749_RevB). Samples were stained using Xenium Multi-Tissue Stain Mix and imaged on Xenium Analyzer instrument according to the manufacturer’s protocol (CG000749_RevB).

We used “Human lung” 289 gene panel from 10x Genomics supplemented with 94 add-on genes (panel ID J2A6YV; **Table S14**). Single-cell spatial transcriptomic data were generated in two independent Xenium runs using the same tissue blocks. Due to the instrument malfunction, the first run (samples with “:1” suffix) was aborted mid-run, and slides were left on the instrument for ∼48 hours. The slides were recovered, stored at 4°C for an additional 24 hours, and then the run was successfully repeated using a new set of reagents. To assess the impact of technical issues on the data quality, we have prepared additional sections from the same tissue blocks (samples with “:2” suffix) and analyzed them using the same gene panel and tissue segmentation kit (replacement reagents were kindly provided by 10x Genomics). The second run completed without technical issues and produced comparable data quality between two runs (**Figure S5c**).

### Single-cell spatial transcriptomic analysis

Cell segmentation was performed with 10X Xenium onboard cell segmentation (Xenium Ranger v 2.0.0.10, 10x Genomics). For each sample, Xenium generated an output file of transcript information including x and y coordinates, corresponding gene target, assigned cell, a binary flag for whether the transcript was expressed over a nucleus, and quality score. Low-quality transcripts (qv < 20) and transcripts corresponding to blank/negative probes were removed.

Sample SSc 2•1 was from the right upper lobe from a SSc-ILD patient who had poorly differentiated non-small cell stage IIIB lung cancer in the right upper lobe, which was treated with immunotherapy, chemotherapy, and radiation two years prior to lung transplantation. We used data from this sample for clustering, cell type identification and niche discovery but excluded it from statistical comparison between donor lungs and lungs from patients with SSc-ILD.

Seurat v5^55^ was used to perform further quality filtering and visualization. A single merged Seurat object was created for all samples based on ROI count matrices and metadata files with nuclei coordinates and area. Nuclei were retained according to the following criteria: ≥ 3 unique genes and ≥ 5 total counts. Because Xenium outputs coordinates based on each slide, which results in samples with shared coordinates across multiple slides, the nuclei coordinates were manually adjusted for visualization so that no samples overlapped during plotting. These adjusted nucleus coordinates were added to the Seurat object and h5ad object as meta-data columns. Seurat v5 was used to perform dimensionality reduction, clustering, and visualization. Gene expression was normalized per cell using Seurat’s `scTransform`^56^ function. Dimensionality reduction was performed with PCA on 383 HVG. Cells were clustered based on this dimensionality reduction using the Louvain algorithm, using 40 PCs as input. Uniform Manifold Approximation and Projection (UMAP) plots of the data were generated using the same number of PCs as used for PCA. Cells were annotated using a combination of marker genes, generated using Seurat v5’s `FindMarkers` function, and spatial information, including cell morphology and position. The CellCharter^57^ package in Python was used to partition cells into 11 spatial niches using k-means clustering based on the transcriptomic expression of the nearest 6 neighboring cells. Spatial abundance analysis was performed by summing cells per cell type per sample. These per sample fractions were then compared across cell types using Welch’s paired t-test with FDR correction.

### Alveolar cell type composition analysis

We manually added 4–9 square regions to all 6 donor sections. Inside each region we manually drew the borders of alveolar spaces fully or almost fully contained within the regions in Xenium explorer software, and exported the coordinates of the polygons for each alveolus in csv format. Next, we loaded all csv files with alveoli polygon coordinates in Python, and identified cells belonging to each alveoli using centroid cell coordinates from Xenium output. For alveolar macrophages and alveolar epithelial cells, we assigned them to the nearest alveolus if their centroids were within 10μm of the alveolar polygon. To normalize for alveoli size, we computed alveoli areas and divided the cell counts by their corresponding alveoli areas and reported cell type abundance per 10,000μm alveolar area. For cytokine expression association analysis, we separated alveoli into cytokine-expressing (at least 1 count of the cytokine mRNA detected in cells of interest) and cytokine non-expressing, and computed average cell type abundance per area for each slide, then compared the paired observations using Wilcoxon signed-rank test.

### Reanalysis of publicly available datasets

To enable analysis of drug target gene expression across different datasets we reanalyzed data from Gao et al. (GSE159354)^15^, Valenzi et al. (GSE128169)^12^, and Papazoglou et al. (GSE212109)^36^. Integration and clustering were performed using scvi-tools and scanpy as described above. Clusters of low-quality cells were removed. We manually annotated cluster cell types. Analysis of expression of selected genes of interest was performed using a pseudobulk approach on a per-cluster level as described above.

### Identification of drugs that are in clinical trials for SSc-ILD

We performed a search of the ClinicalTrials.gov database on May 1st, 2025, using the search term “Systemic Sclerosis Associated Interstitial Lung Disease”.

## Data availability

Single-cell RNA-seq counts tables and integrated objects are available through the Gene Expression Omnibus with accession numbers GSE303858 and GSE303859. We also used publicly available single-cell RNA-seq data from GSE15935415, GSE12816912, and GSE212109. Xenium counts tables and Xenium Explorer bundles are available with accession number GSE303048

## Code availability

All code used for analysis is available at https://github.com/NUPulmonary/2025_Markov_SSc. Single-cell RNA-seq and single-cell spatial data can be explored via data browsers at https://sqlifts.fsm.northwestern.edu/public/Markov_2025_SSc/.

## Acknowledgements

We thank all patients and volunteers who provided their samples and data for this study. This research was supported in part through a generous gift from Kimberly Querrey and Louis A. Simpson. This research was supported by the Simpson Querrey Lung Institute for Translational Science (SQLIFTS), Northwestern University. This research was also supported by the computational resources and staff contributions provided for the Quest high-performance computing facility at Northwestern University, which is jointly supported by the Office of the Provost, the Office for Research and Northwestern University Information Technology. This research was also supported in part through the computational resources and staff contributions provided by the Genomics Compute Cluster, which is jointly supported by the Feinberg School of Medicine, the Center for Genetic Medicine and Feinberg’s Department of Biochemistry and Molecular Genetics, the Office of the Provost, the Office for Research and Northwestern Information Technology. The Genomics Compute Cluster is part of Quest, Northwestern University’s high-performance computing facility, with the purpose of advancing research in genomics. We thank Jackie Milhans, Alper Kinaci, Scott Coughlin, and all members of the Research Computing and Data Services team at Northwestern for their support. We thank Drs. Liza Hazelwood and Lauren Reinke-Breen for productive discussions of the study results. This work is dedicated to the benefit of all sentient beings. Northwestern University Flow Cytometry Core Facility is supported by the National Cancer Institute Cancer Center support grant (no. P30 CA060553) awarded to the Robert H. Lurie Comprehensive Cancer Center. Cell sorting was performed on a BD FACS Aria SORP cell sorter purchased with the support of the National Institutes of Health (NIH, grant no. 1S10OD011996-01). Integrative genomic services were performed by the Metabolomics Core Facility at Robert H. Lurie Comprehensive Cancer Center of Northwestern University. Next-generation sequencing was performed with support from the Simpson Querrey Institute for Epigenetics.

N.S.M. was supported by the American Heart Association (grant no. 24PRE1196998 https://doi.org/10.58275/AHA.24PRE1196998.pc.gr.190609).

A.J.E. was supported by the NIH (grant no. L30HL149048), and Pulmonary Fibrosis Foundation Scholars Award.

P.A.R. was supported by NIH (grant no. K08HL146943), the Parker B. Francis Fellowship, and the ATS Foundation/Boehringer Ingelheim Pharmaceuticals Inc. Research Fellowship in IPF.

A.B. was supported by the NIH (grant nos. P01HL169188, R01HL147290, R01HL145478 and R01HL147575).

R.M.T. was supported by the NIH (grant nos. R01ES034350 and R01ES027574).

H.P. was supported by the NIH (grant nos. R01AR080513, P01HL169188).

A.P.L. was supported by the NIH (grant no. R01HL134800).

G.R.S.B. was supported by Simpson Querrey Lung Institute for Translational Science, the NIH (grant nos. P01AG049665, P01HL154998, U54AG079754, R01HL147575, R01HL158139, R01HL147290,

R21AG075423 and U19AI135964) and the Veterans Administration (award no. I01CX001777).

C.J.G. was supported by NIH (grants nos. GM129312, HL163611, and HL134800), and a grant from Boehringer Ingelheim IIS2017-10679.

A.V.M. was supported by the NIH (grant nos. U19AI135964, P01AG049665, P01HL154998, P01HL169188, U19AI181102, R01HL153312, R01HL158139, and R01ES034350), and research grants from AbbVie and Merck.

M.E.H. was supported by the NIH (grant no. AR073270), and research grants from Boehringer Ingelheim.

The funders had no role in the study design, data collection and analysis, decision to publish, or manuscript preparation.

---

**Supplementary Figure 1:**
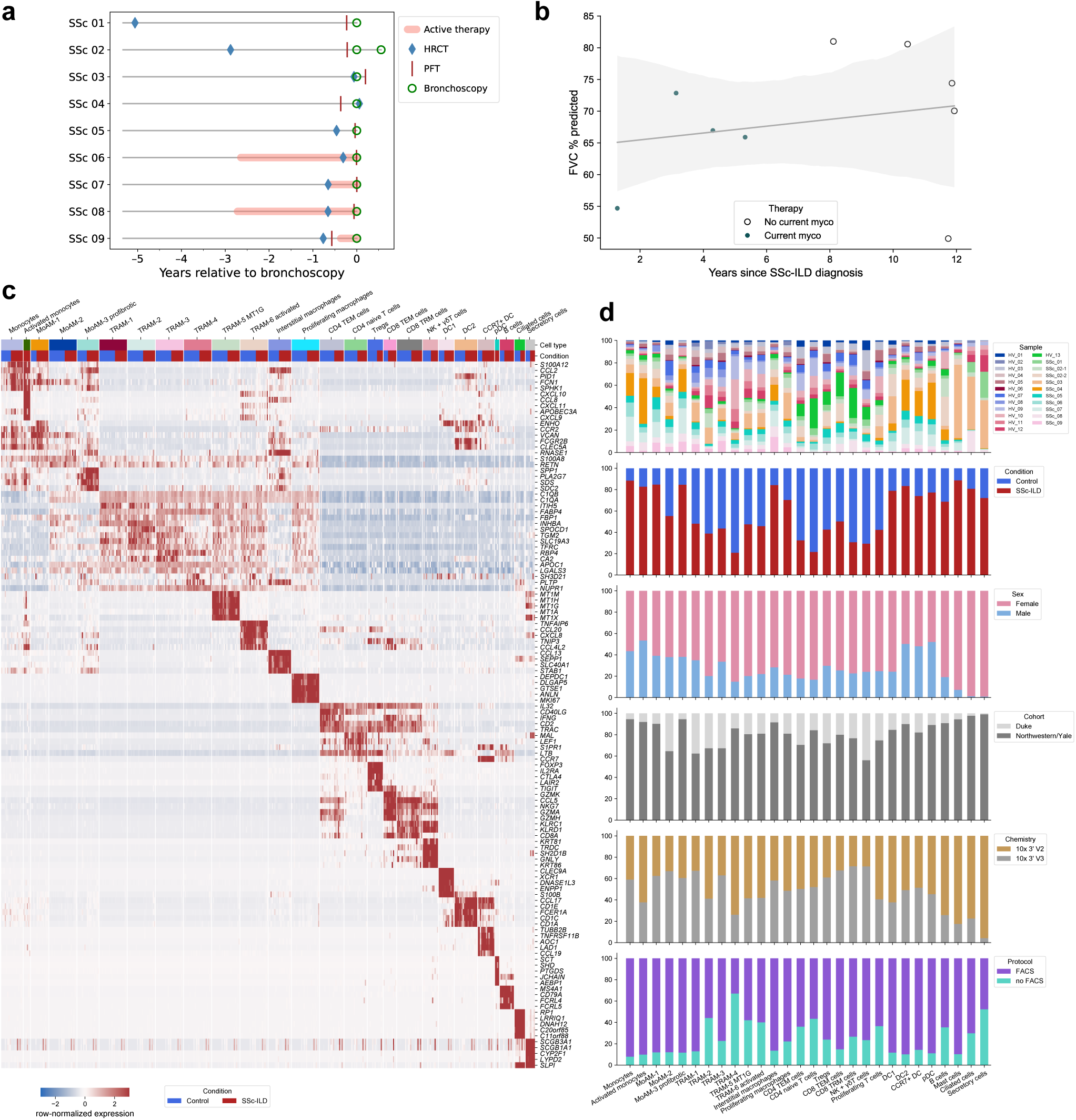
Characterization of patients with SSc-ILD and scRNA-seq analysis of cells isolated from BALF from controls and patients. **a**. Schematic representation of the clinical course and selected diagnostic tests and interventions in 9 patients with SSc-ILD preceding the bronchoscopy. Timing of CT scans of the chest, bronchoscopy, pulmonary function testing (PFT) and treatment with mycophenolate mofetil are annotated as symbols. **b**. Correlation between pulmonary function (% of predicted forced vital capacity, FVC) and duration of SSc-ILD (years). Red and blue symbols indicate patients on active therapy with mycophenolate or not treated with mycophenolate, respectively. **c**. Heatmap showing expression of the genes used as markers to identify cell types in the integrated single-cell RNA-seq object from **Figure 1a**. **d**. Barplots illustrating cluster composition by sample, condition, sex, cohort and chemistry.

**Supplementary Figure 2:**
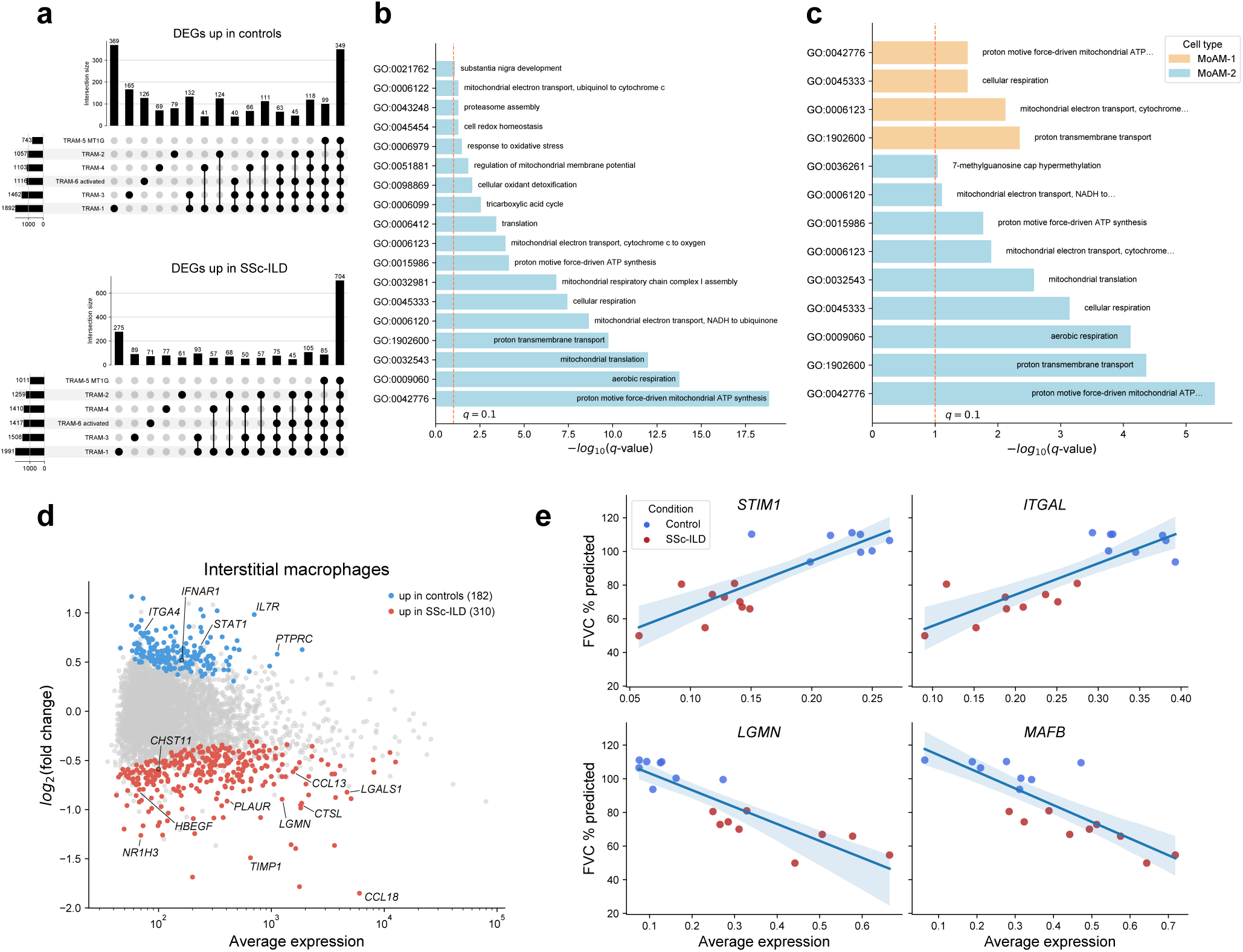
Macrophage and T cell gene programs associate with SSc-ILD. **a**. UpSet plots illustrating unique and shared DEGs in TRAM subsets, only sets with at least 40 genes are shown. **b**. Barplot of significantly enriched (*q*-value < 0.1) GO processes among the set of common DEGs upregulated in TRAM subsets in patients with SSc-ILD. **c**. Barplot of significantly enriched (*q*-value < 0.1) GO processes for DEGs in MoAM-1 and MoAM-2 subsets upregulated in patients with SSc-ILD. **d**. Differentially expressed genes in interstitial macrophages between controls and patients. Significant genes (*q*-value < 0.05) are highlighted in color. Selected genes of interest are labeled. **e**. Correlation analysis between average expression of *STIM1*, *ITGAL*, *LGMN*, and *MAFB* and % predicted FVC. Pearson correlation, shaded area corresponds to the 95% confidence interval.

**Supplementary Figure 3:**
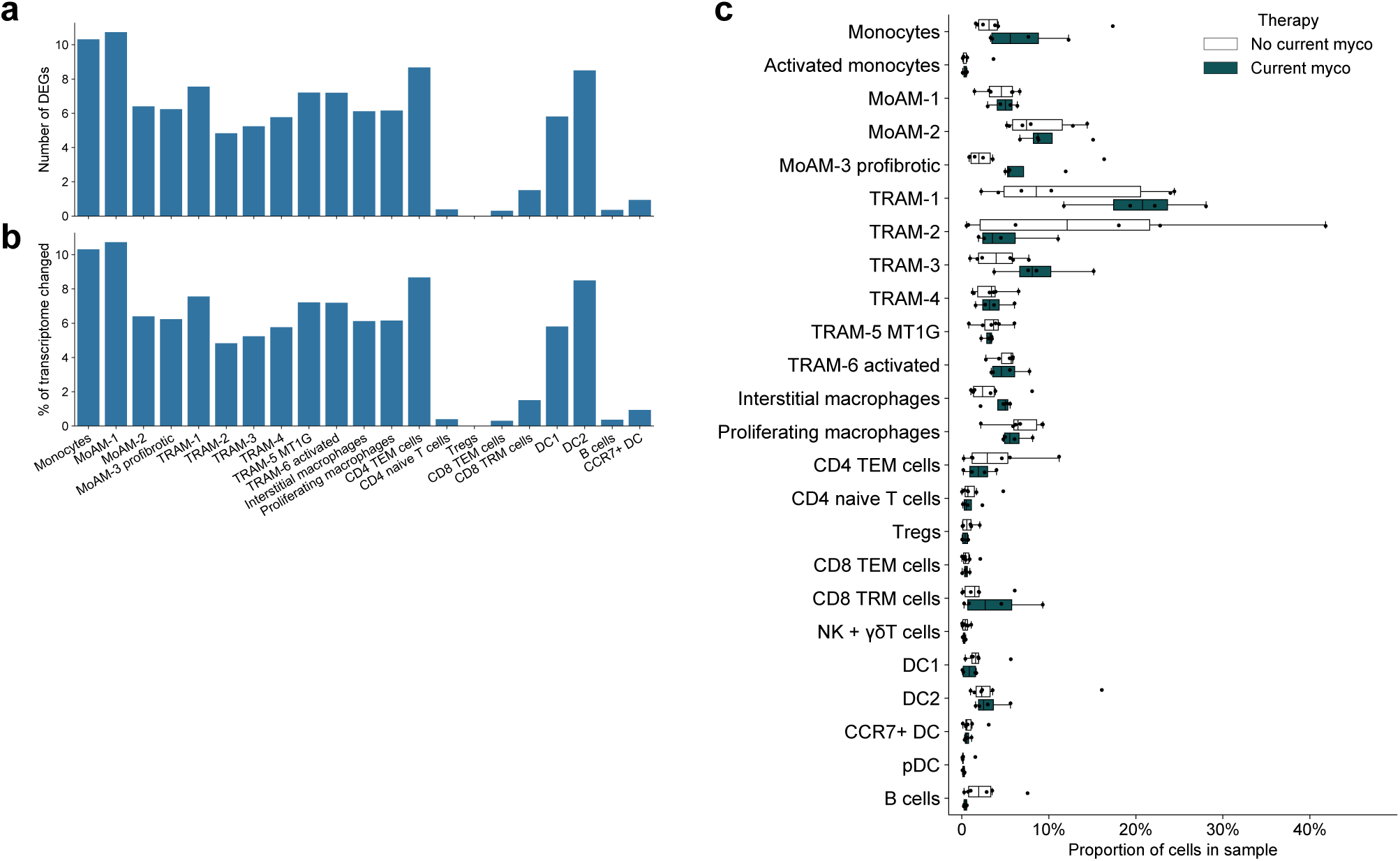
Mycophenolate therapy is associated with transcriptomic changes in immune cells. **a**. Number of differentially expressed genes (DEGs, *q*-value < 0.05) in different cell types between patients receiving or not receiving active mycophenolate therapy. **b**. Percent of transcriptome within different cell types represented by DEGs between patients receiving or not receiving active mycophenolate therapy. **c**. Proportions of cell clusters represented in the UMAP in **Figure 1a** between patients on active therapy with mycophenolate in comparison to untreated patients. *q*-value < 0.05 are shown next to pairs of boxplots (Mann-Whitney U tests with FDR correction).

**Supplementary Figure 4:**
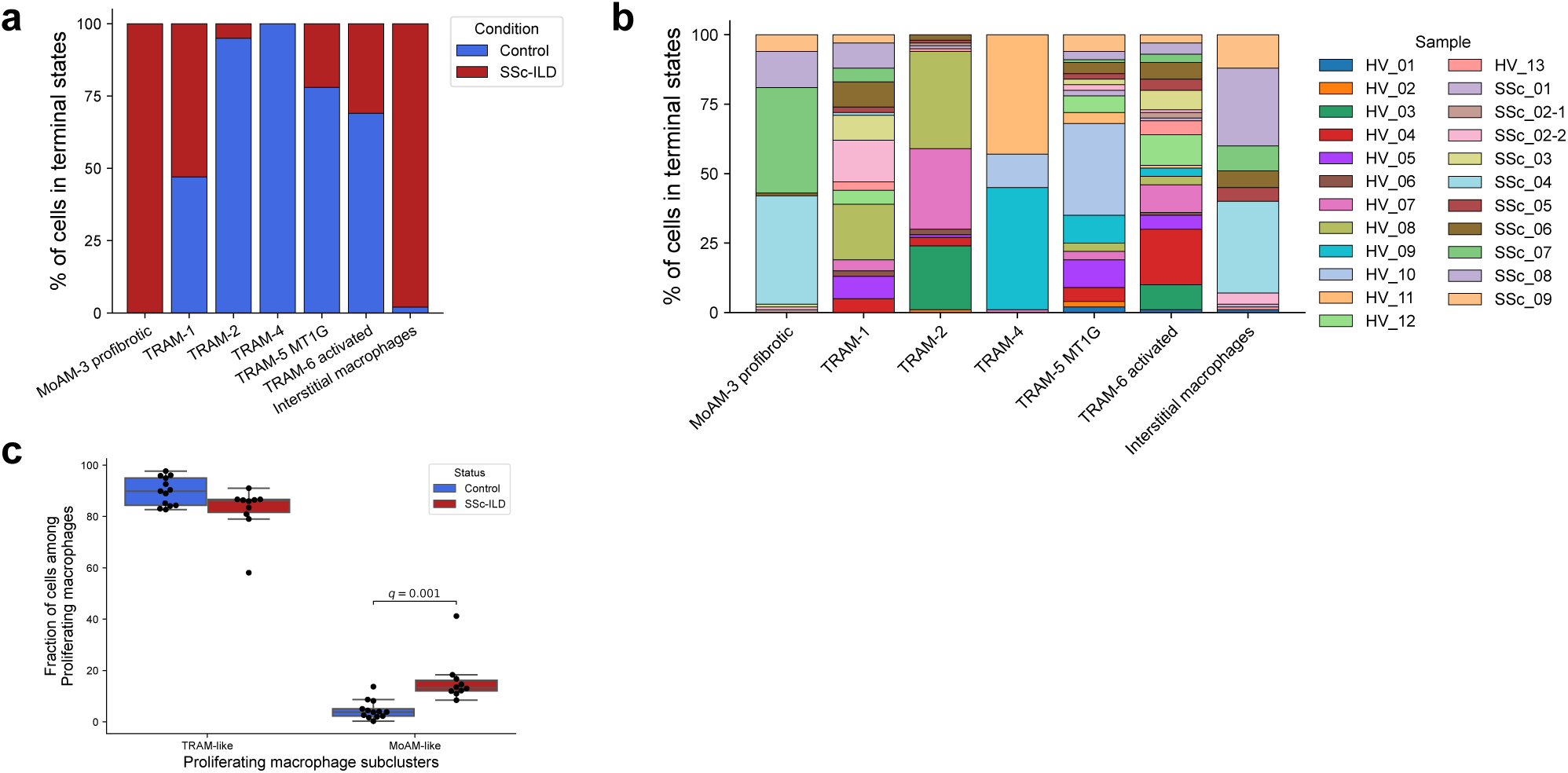
Profibrotic *SPP1+* MoAM represent a pathologic monocyte-to-macrophage differentiation trajectory in SSc-ILD. **a**. Composition of cells in terminal states based on disease status. **b**. Composition of cells in terminal states based on sample. **c**. Proportions of TRAM- and MoAM-like macrophages among proliferating macrophages from **Figure 1a**. *q*-value < 0.05 are shown next to pairs of boxplots (Mann-Whitney U tests with FDR correction).

**Supplementary Figure 5:**
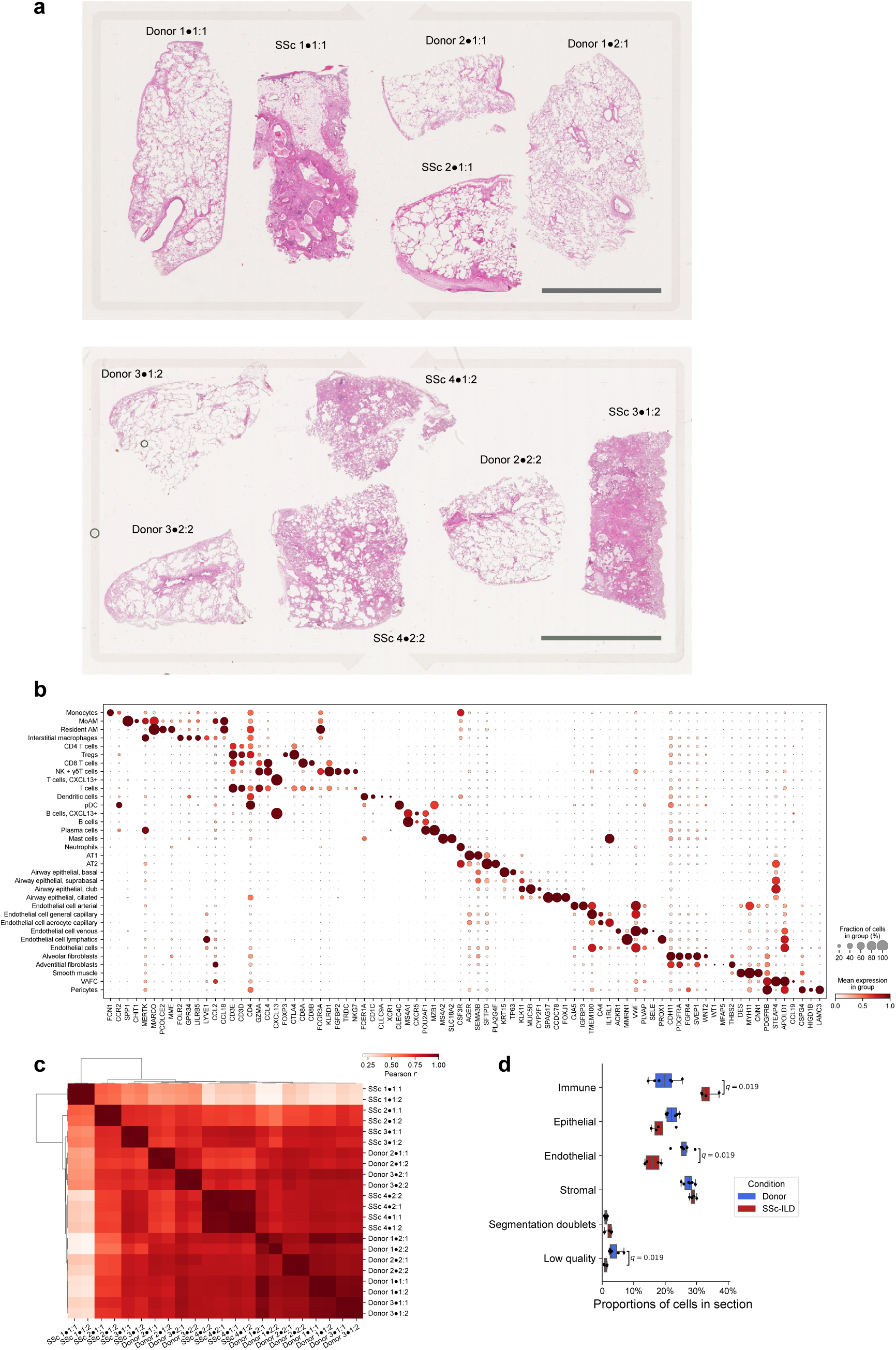
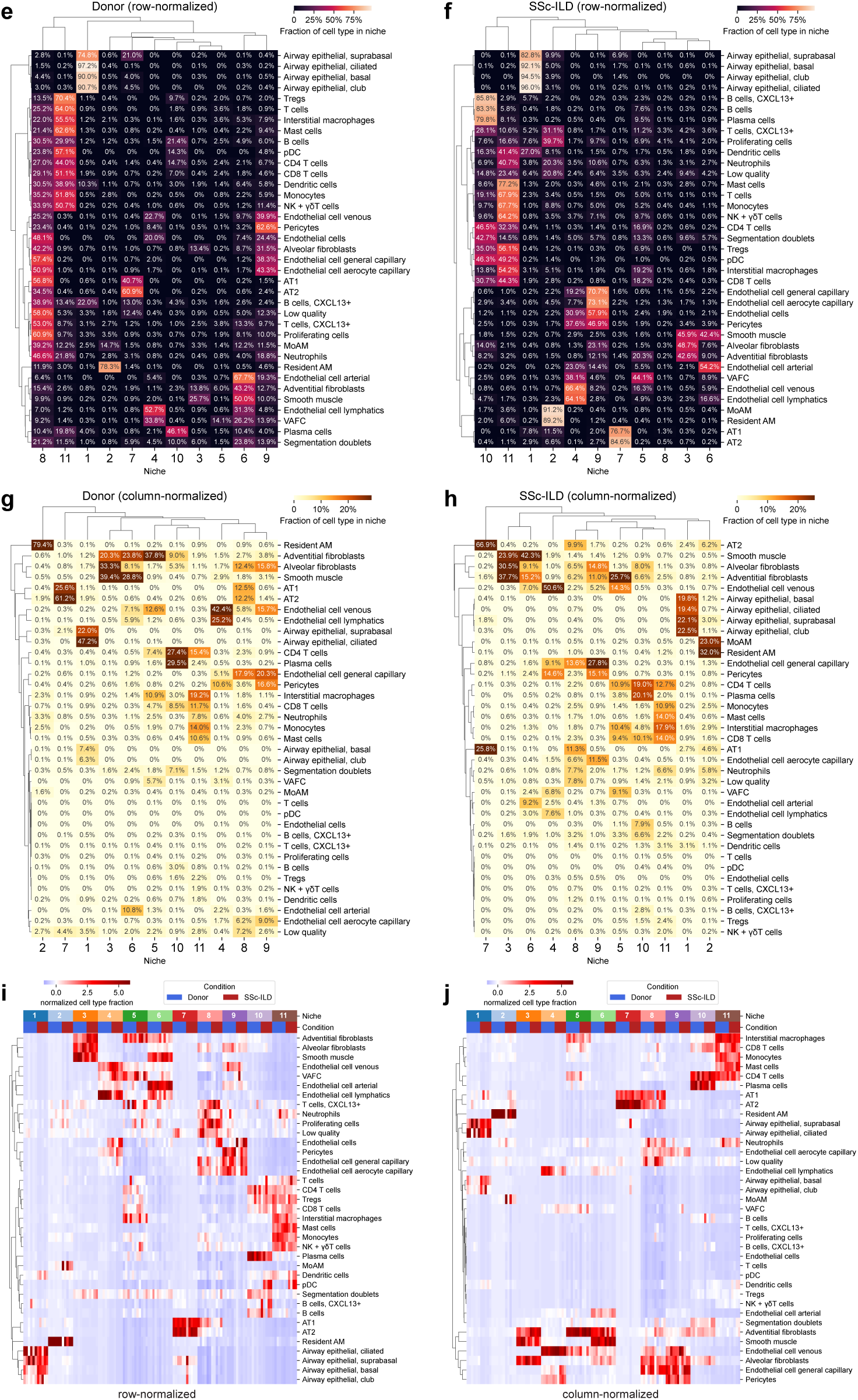
Single-cell spatial profiling identifies niche for profibrotic *SPP1+* MoAM. **a**. Histology of the lungs from patients with SSc-ILD and donor lungs from spatial transcriptomic profiling. Scale bar 5 mm. Donor or patient, tissue sample and slide number are indicated (see Table 3). **b**. Heatmap showing expression of the genes used as markers to identify cell types in the integrated single-cell spatial object from Figure 5a. **c**. Hierarchical clustering of Pearson correlation coefficients between cell type numbers for each slide at the most granular cell type annotation (level 4 in **Table S9**). Average linkage method over cosine distances. **d**. Comparison of proportions of main cell lineages (level 1 in **Table S9**) between lungs from donors and patients with SSc-ILD. *q*-value < 0.05 are shown above each pair of boxplots, Mann-Whitney U tests with FDR correction. **(e**. Hierarchical of niche composition by cell type (level 3 in **Table S9**) in donor lungs. Data scaled across rows. Numbers indicate % of cell type belonging to a specific niche. Ward linkage method over euclidean distances. **f**. Hierarchical clustering of niche composition by cell type (level 3 in **Table S9**) in lungs from patients with SSc-ILD. Data scaled across rows. Numbers indicate % of cell type belonging to a specific niche. Ward linkage method over Euclidean distances. **g**. Hierarchical clustering of niche composition by cell type (level 3 in **Table S9**) in donor lungs. Data scaled across columns. Numbers indicate % of niche occupied by a specific cell type. Ward linkage method over Euclidean distances. **h**. Hierarchical clustering of niche composition by cell type (level 3 in **Table S9**) in lungs from patients with SSc-ILD. Data z-scaled across columns. Numbers indicate % of niche occupied by a specific cell type. Ward linkage method over Euclidean distances. **i**. Hierarchical clustering of niche composition by cell type (level 3 in **Table S9**) in lungs from donors and patients with SSc-ILD. Data z-scaled across rows. Each column represents a single sample. Ward linkage method over Euclidean distances. **j**. Hierarchical clustering of niche composition by cell type (level 3 in **Table S9**) in lungs from donors and patients with SSc-ILD. Data z-scaled across columns. Each column represents a single sample. Ward linkage method over Euclidean distances.

**Supplementary Figure 6:**
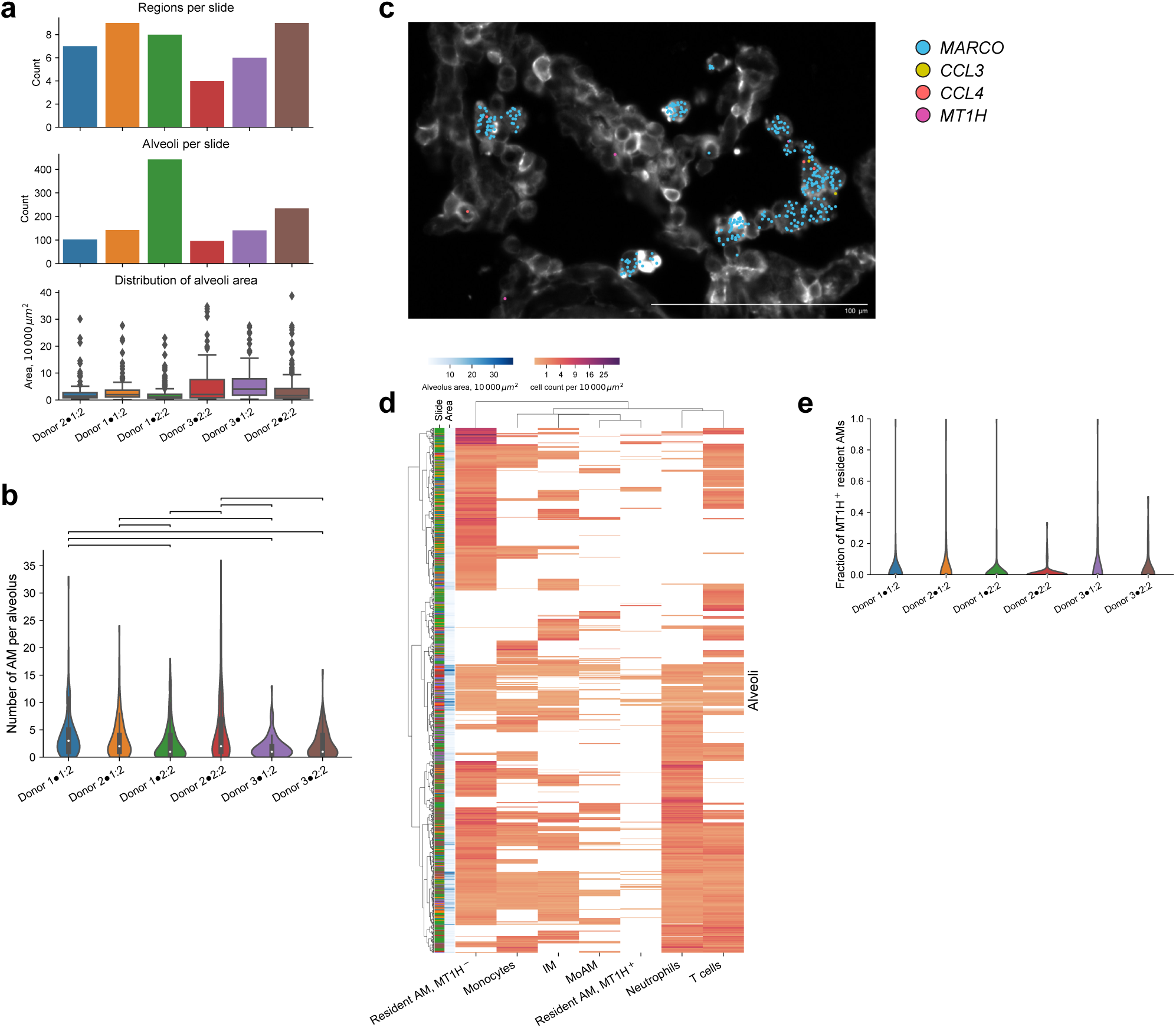
Transcriptionally distinct alveolar macrophage subsets are co-localized within the same alveoli. **a**. Number of regions (top) and alveoli (middle) manually annotated per slide from the donor lungs, and the distribution of alveoli size (bottom). **b**. Distribution of the number of alveolar macrophages detected per single alveolus across donor samples. Mann-Whitney U tests with FDR correction. *q*-value < 0.05 are shown as brackets. **c**. Representative Xenium image showing co-localization of MARCO+ alveolar macrophages positive and negative for *CCL3*, *CCL4*, and *MT1H* within the same alveolus in the donor lungs. Cell boundary stain from Xenium cell segmentation kit is shown. Each dot represents corresponding transcript (see legend). **d**. Hierarchical clustering on abundance of immune cell composition in the alveolar spaces in the donor lungs shows that *MT1H+* TRAM coexist together with *MT1H–* TRAM. Ward linkage over Euclidean distances. **e**. Violin plot of fractions of *MT1H+* TRAM out of all TRAM per alveolus across donor samples. White dot indicated median value.

**Supplementary Figure 7:**
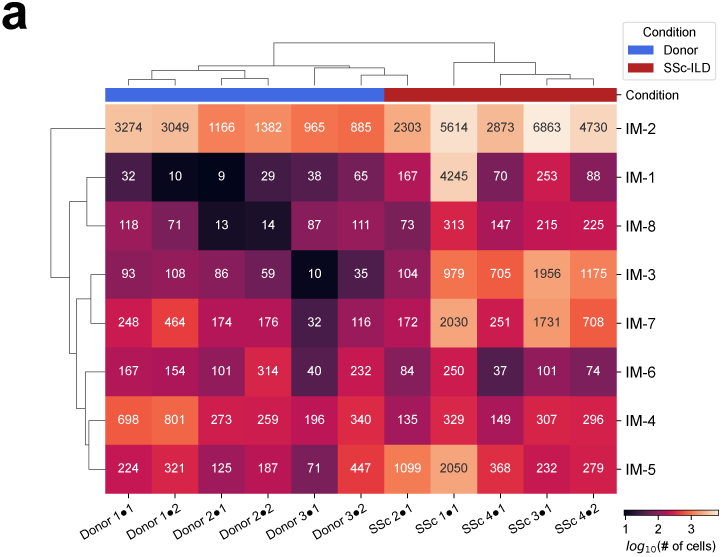
Interstitial macrophages “spill” into the alveolar space. **a**. Heatmap of absolute cell count for interstitial macrophage subsets per section. Ward linkage over Euclidean distances in log_10_ space.

**Supplementary Figure 8:**
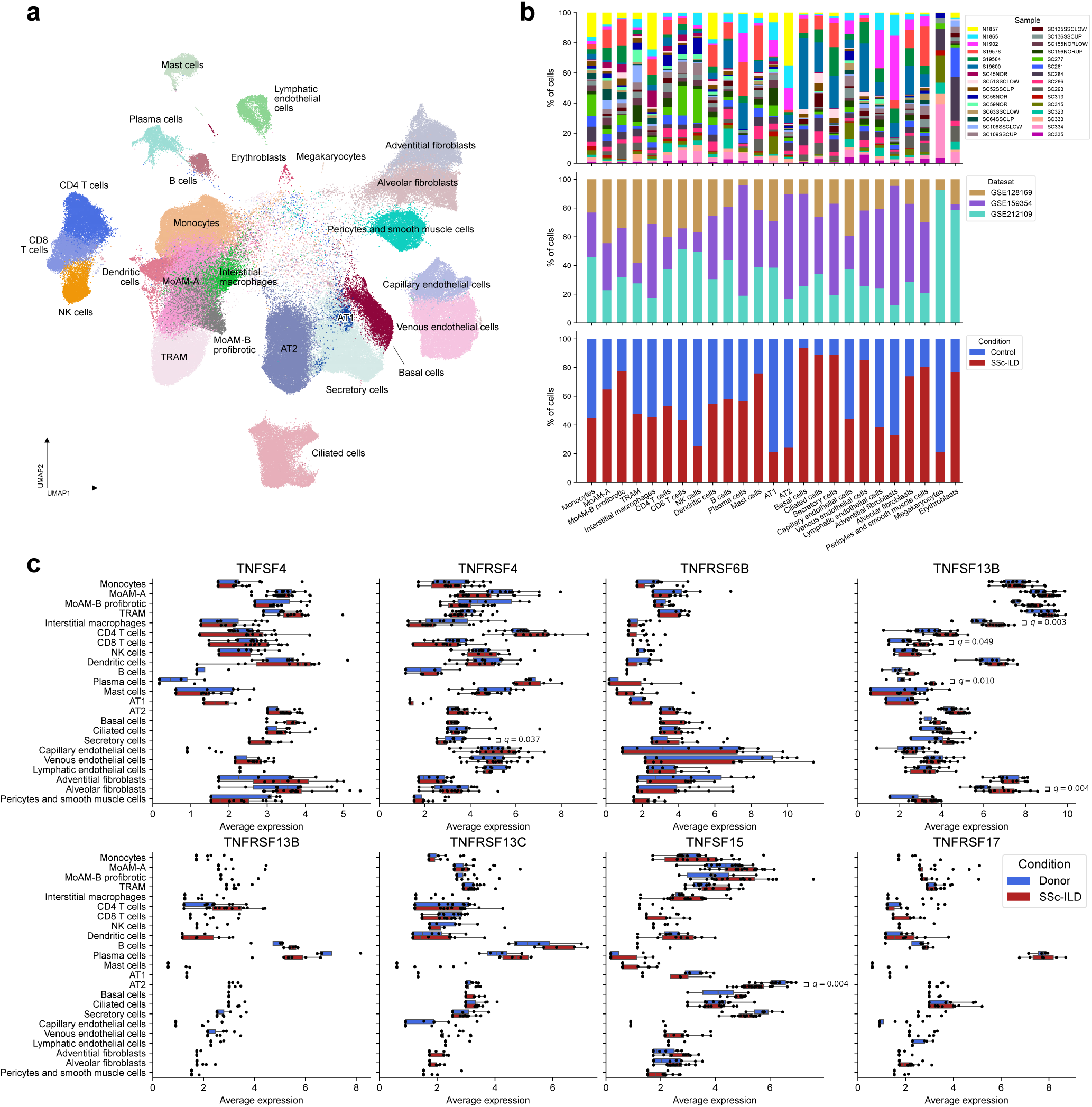
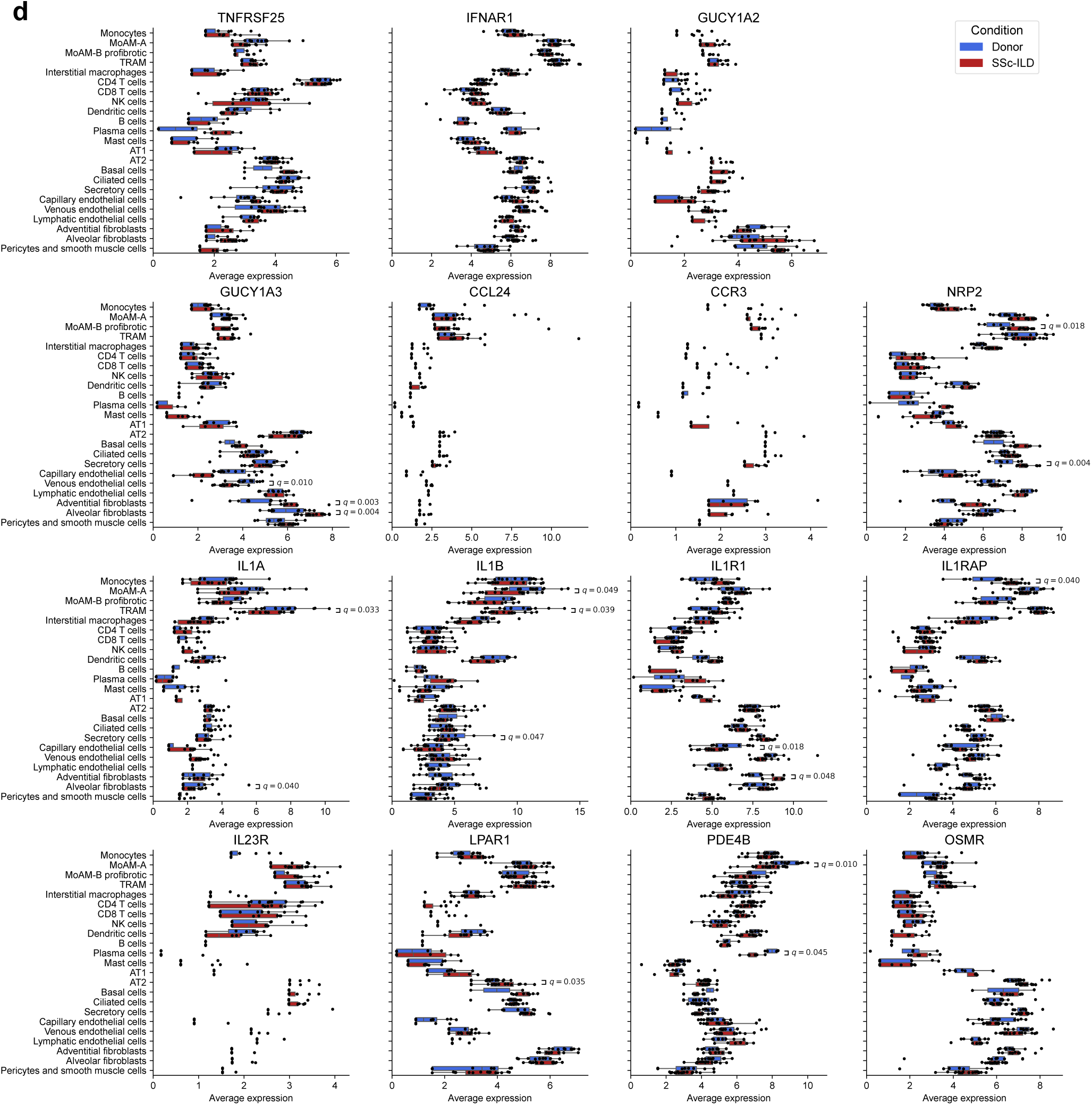
Pharmacological targets in patients with SSc-ILD. **a**. UMAP of the integrated object from Gao et al. (GSE159354), Valenzi et al. (GSE128169), and Papazoglou et al. (GSE212109). Clusters are annotated according to the Human Lung Cell Atlas labels. **b**. Barplot illustrating cluster composition by sample, study and condition (donor vs SSc-ILD). **c**. Expression of genes encoding ligands, receptors, or enzymes that are currently in clinical trials for patients with SSc-ILD. Reanalysis of publicly available data from patients with end-stage SSc-ILD. *q*-value < 0.05 are shown above each pair of boxplots, DESeq2 tests with FDR correction.

## List of supplementary tables

Supplementary Table S1: Marker genes for scRNA-seq object, **Figure 1a**.

Supplementary Table S2: DEGs between controls and patients. **Figure 2a**.

Supplementary Table S3: GSEA on DEGs between controls and patients. **Figure 2c**.

Supplementary Table S4: GO biological processes for DEGs in TRAM subsets between controls and patients. **Figure S2b**.

Supplementary Table S5: GO biological processes for DEGs in MoAM subsets between controls and patients. **Figure S2c**.

Supplementary Table S6: Biomarker candidate genes. **Figure S2e**.

Supplementary Table S7: DEGs between patients on active therapy with mycophenolate in comparison to untreated patients. **Figure 3**.

Supplementary Table S8: GSEA on gene lists between patients on active therapy with mycophenolate in comparison to untreated patients. **Figure 3a**.

Supplementary Table S9: Hierarchical cell type annotation for single-cell spatial dataset. **Figure 5**.

Supplementary Table S10: Niche composition on a per-section level, normalized by cell type and by niche.

Supplementary Table S11: Drugs that are in clinical trials for SSc-ILD.

Supplementary Table S12: Marker genes for integrated single cell RNA-seq object from patients with end-stage SSc-ILD.

Supplementary Table S13: DEGs for targets of drugs in table S11 in scRNA-seq object from patients with end-stage SSc-ILD.

Supplementary Table S14: Human lung Xenium panel with 94 add-on genes.

